# Skeletal Muscle Satellite Cells Co-Opt the Tenogenic Gene *Scleraxis* to Instruct Regeneration

**DOI:** 10.1101/2023.12.10.570982

**Authors:** Yun Bai, Tyler Harvey, Colin Bilyou, Minjie Hu, Chen-Ming Fan

## Abstract

Skeletal muscles connect bones and tendons for locomotion and posture. Understanding the regenerative processes of muscle, bone and tendon is of importance to basic research and clinical applications. Despite their interconnections, distinct transcription factors have been reported to orchestrate each tissue’s developmental and regenerative processes. Here we show that *Scx* expression is not detectable in adult muscle stem cells (also known as satellite cells, SCs) during quiescence. *Scx* expression begins in activated SCs and continues throughout regenerative myogenesis after injury. By SC-specific *Scx* gene inactivation (ScxcKO), we show that *Scx* function is required for SC expansion/renewal and robust new myofiber formation after injury. We combined single-cell RNA-sequencing and CUT&RUN to identify direct Scx target genes during muscle regeneration. These target genes help explain the muscle regeneration defects of ScxcKO, and are not overlapping with *Scx*-target genes identified in tendon development. Together with a recent finding of a subpopulation of *Scx*-expressing connective tissue fibroblasts with myogenic potential during early embryogenesis, we propose that regenerative and developmental myogenesis co-opt the *Scx* gene via different mechanisms.

## Introduction

Regeneration of adult skeletal muscles following injury is initiated by the activation and proliferation of satellite cells (SCs). After extensive proliferation, progenitors undergo differentiation and fusion with each other or existing myofibers to recreate functional muscle tissue(Yin, Price et al. 2013, Liu, Nelson et al. 2014, Fukada, Higashimoto et al. 2022). The intrinsic and extrinsic factors regulating myogenesis have been extensively investigated. The key transcription factors governing this process are largely the same as those deployed during embryogenesis, including paired-homeodomain proteins Pax3 and Pax7, basic helix-loop-helix (bHLH) myogenic regulatory factors (MRFs) such as Myf5 and MyoD1, and the myocyte enhancer factor 2 (MEF2) family; however, their relative contribution or redundancy vary between the two processes(Hernandez-Hernandez, Garcia-Gonzalez et al. 2017). To date, resident Pax7^+^ SCs are recognized as the major source of muscle stem cells in adult limb muscles. None of these myogenic transcription factors are known to participate in tendon development or regeneration.

Both muscle and tendon progenitors reside in the somite during embryonic development but are located in different compartments. *Pax3* and *Pax7* are expressed in the dermomyotome, which gives rise to the myotome expressing *Myf5* and/or *MyoD1*. The syndetome, on the other hand, is defined by the expression of the earliest tenogenic progenitor marker *Scx*, and gives rise to tendon and ligament(Brent, Schweitzer et al. 2003). Like Myf5 and MyoD1, Scx is a bHLH transcription factor, and they all bind to a DNA sequence motif called the E-box(Cserjesi 1995). *Scx* expression persists in mature tenocytes, ligaments, and connective tissue fibroblasts (CT)(Murchison, Price et al. 2007). *Scx* mutant mice have poorly developed tendons with drastically reduced expression of tendon matrix genes(Murchison, Price et al. 2007, Yoshimoto, Takimoto et al. 2017, Shukunami, Takimoto et al. 2018). In adult tendon regeneration, the Tppp3^+^Pdgfra^+^ tendon stem cell population turn on *Scx* for tendon regeneration(Harvey, Flamenco et al. 2019). Lastly, *Scx* function is required in post-natal tendon growth and regeneration(Howell, Chien et al. 2017, Sakabe, Sakai et al. 2018, Gumucio, Schonk et al. 2020, Korcari, Muscat et al. 2022).

Intriguingly, lineage tracing using a constitutive *Scx^Cre^* in mouse embryos found descendant cells in cartilage, tendon, ligament, muscle, and muscle interstitial CT(Yoshimoto, Takimoto et al. 2017, Esteves de Lima, Blavet et al. 2021, Ono, Schlesinger et al. 2023), suggesting that *Scx* is expressed either in several distinct musculoskeletal subpopulations, or in a common progenitor that gives rise to different fates. Ablation of embryonic Scx^+^ cells cause a change in muscle bundling(Ono, Schlesinger et al. 2023), presumably due to the loss of instructive cues from the tendon (or CT) to form proper muscle pattern(Kardon 1998). In adult muscles, Hic1^+^ quiescent mesenchymal progenitors (MPs) give rise to Scx^+^ cells in the muscle interstitial compartment, and ablation of Hic1^+^ cells negatively impacts muscle regeneration(Scott, Arostegui et al. 2019). Muscle interstitial Scx^+^ cells engrafted into the muscle contribute only to extracellular matrix remodeling(Giordani, He et al. 2019). A survey of muscle interstitial CT assigned a sub-population of cells expressing tendon markers (including *Scx*) as paramysial cells - cells lining next to the perimysium that wraps around muscle fascicles(Muhl, Genove et al. 2020). Furthermore, Strenzke and colleagues showed that secretome from Scx overexpressed cells could significantly increase myoblast fusion and metabolic activity *in vitro*(Strenzke, Alberton et al. 2020). Collectively, these data indicate that while some embryonic Scx^+^ cells can incorporate into myofibers, adult Scx^+^ cells contribute to skeletal muscle architecture and repair/regenerative process in a paracrine manner.

Serendipitously, in the ScxGFP transgenic mouse Tg-ScxGFP(Pryce, Brent et al. 2007), we observed GFP fluorescence in SCs and regenerating myofibers after injury. We conducted a series of experiments to show that endogenous *Scx* is expressed in activated SC after injury. We show that *Scx* is functionally relevant in muscle regeneration by inactivating *Scx* in *Pax7^+^* SCs (ScxcKO).We employed single-cell RNA-sequencing (scRNA-seq) and CUT&RUN to define Scx’s target genes during muscle differentiation and fusion. Down-regulation of Scx’s target genes such as *Mef2a*, *Cflar*, *Capn2*, and *Myh9,* explains the regenerative defects of ScxcKO mice. In contrast to adult Scx^+^ muscle CT and embryonic muscle-forming Scx^+^ cells, our findings reveal a previously unappreciated role of *Scx* in adult Pax7^+^ SCs.

## Results

### ScxGFP transgene is expressed in the regenerative myogenic lineage

When we analyzed tibialis anterior (TA) muscles of the Tg-ScxGFP (ScxGFP) mice, scattered GFP^+^ cells were found in the interstitial space, but not in quiescent Pax7^+^ SCs nor in myofibers (Fig. S1A). Unexpectedly, we found GFP signal in injured muscles. In cardiotoxin (CTX) injured TA muscles of ScxGFP mice at 5 days post-injury (dpi) (Fig.1A), we found GFP colocalized with Pax7^+^ SCs (Fig.1B). GFP also overlaps with committed myogenic progenitor marker MyoD1, myocyte marker Myogenin (Myog), and myosin heavy chain (MHC) in terminally differentiated myofibers (Fig. 1C; Fig. S1B-D for split channels). When we analyzed muscles administered with 5-ethynyl-2′-deoxyuridine (EdU) for 5 days after CTX injury (Fig. 1A), GFP was found to colocalize with proliferated (EdU^+^) Pax7^+^ cells (Fig. 1D). Thus, ScxGFP is expressed, albeit at varying levels, in the myogenic lineage during regenerative process.

**Figure 1:**
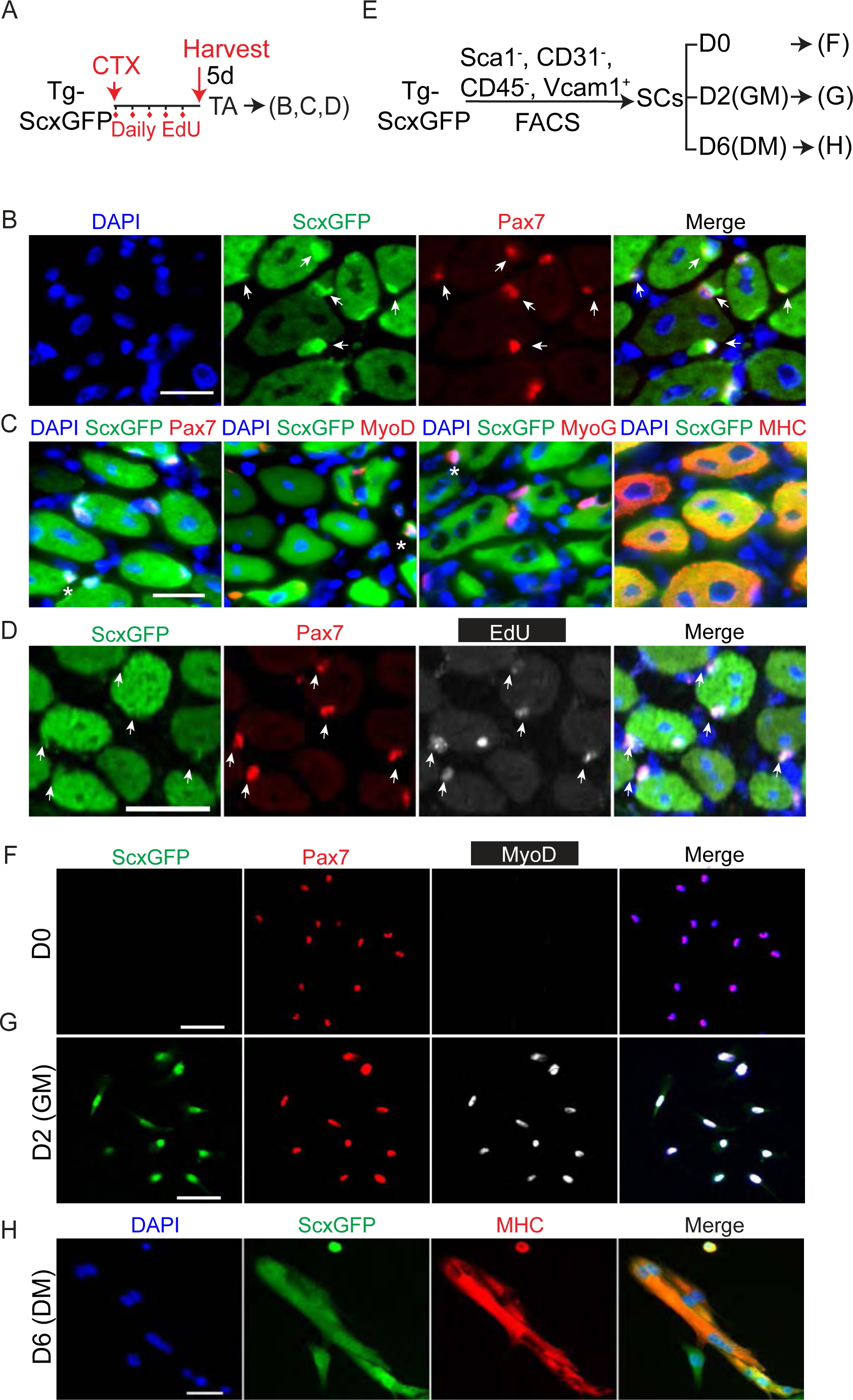
Adult regenerative myogenic cells express the transgene ScxGFP. A. Experimental scheme for data in (B-D). Tg-ScxGFP (ScxGFP) mice were injured by CTX to the TA muscle, followed by daily EdU administration for 5 days (5d), and their TA muscles were harvested for analysis at 5d post-injury (dpi). B. Muscle samples obtained in (A) were sectioned and stained with Pax7 and GFP (for ScxGFP expression) antibody. Arrows indicate Pax7 and ScxGFP double positive cells; 97.77% Pax7^+^ SCs were ScxGFP^+^ (N = 4 mice; n = 1274 cells). C. Muscle samples obtained in (A) were sectioned and stained in pairs of GFP/Pax7, GFP/MyoD1, GFP/MyoG, GFP/MYH (N = 4 mice). Asterisks indicate cells double-positive for ScxGFP and each respective myogenic marker. All myofibers are GFP and MHC double positive, thus without additional labeling. D. Muscle samples obtained in (A) were sectioned and stained for Pax7 and GFP, followed by EdU reaction (N = 4 mice). Arrows indicate Pax7, ScxGFP, and EdU triple-positive SCs. E. Experimental scheme of SC isolation from Tg-ScxGFP hindlimb muscles using four surface markers (CD31^-^, CD45^-^, Sca1^-^, Vcam1^+^) by FACS. Isolated SCs were assayed immediately after isolation (D0; data in (F)), after culture in growth media for 2d (D2(GM); data in (G)), or after cultured for 4d in GM followed by 2d in differentiation media (DM) (D6(DM); data in (H)). F-G. D0 (F) and D2 cultured (G) SCs obtained in (E) were stained with for GFP (i.e. ScxGFP), Pax7, and MyoD. At D0, no Pax7^+^ cells were GFP^+^ or MyoD^+^. At D2, 95.3% of Pax7^+^ cells were GFP^+^, whereas 99.2% of MyoD^+^ were GFP^+^. (N = 3 mice; n = 1805 cells at D0; n = 1332 cells at D2). H. D6(DM) cells obtained in (E) were stained for GFP (i.e. ScxGFP) and MHC. 94.58% MHC^+^ were GFP^+^. (N= 3 mice; n = 1539 nuclei in MHC^+^ domain examined). Nuclei were stained with DAPI (blue); Scale bars = 20 μm.

We next determined ScxGFP expression in cultured SCs. For this, we employed a four-surface marker fluorescent activated cell sorting (FACS) scheme (Sca1^-^CD31^-^CD45^-^Vcam1^+^)(Liu, Cheung et al. 2015) to purify SCs from hindlimb muscles of ScxGFP mice (Fig. 1E; S1E, F); ∼ 98% of isolated cells were Pax7^+^ (Fig. S1G, H). While Pax7 was detected in the SC immediately after FACS isolation, neither MyoD nor GFP was detected (Fig. 1F). After two days in culture, most cells were Pax7, MyoD and GFP triple positive (Fig. 1G). After switched to differentiation media for 2 days, GFP signal persisted in MHC^+^ myotubes (Fig. 1H). We therefore conclude that ScxGFP expression is initiated after SC becomes activated and continues into differentiated myofibers *in vivo* and *in vitro*.

### Endogenous *Scx* is expressed in activated SCs

To ensure that the ScxGFP expression observed in adult regenerative myogenesis is not caused by mis-expression due to transgene insertion site, we utilized *Scx^CreERT2^* for tamoxifen (TMX) inducible lineage tracing with a tdTomato (tdT) reporter (*R^tdT^*)(Madisen, Zwingman et al. 2010). Two experimental groups with different tamoxifen (TMX) and injury regimen were designed (Fig. 2A): 1) TMX-induced marking before injury, and 2) TMX-induced marking after injury; muscles were harvested at 14 dpi for analysis. Mice treated with TMX before injury showed little to no tdT^+^Pax7^+^ SCs or tdT^+^ regenerative muscle fibers (identified by centrally located nuclei). By contrast, mice treated with TMX after injury showed ∼ 30% of Pax7^+^ SCs and all regenerated myofibers as tdT^+^ at 14 dpi (Fig. 2C-D). Pax7^+^ SC densities were not different between these two groups (Fig. S2A). These data extend the ScxGFP results in that 1) marked interstitial Scx^+^ cells prior to injury do not possess myogenic potential, 2) endogenous *Scx* is expressed in activated SCs for regenerative myogenic lineage-marking, and 3) lineage-marked *Scx*^+^ SCs are capable of renewal as Pax7^+^ SCs at 14 dpi. Examination of two published SC bulk RNA-seq data confirmed *Scx* expression in SCs isolated from wild type(Li, Rozo et al. 2019) and *mdx* mice(Madaro, Torcinaro et al. 2019) (Fig. S2B). Re-analysis of published scRNA-seq data sets of regenerative myogenic cells also uncovered a wide-spread *Scx* expression at 2 dpi (De Micheli, Laurilliard et al. 2020) and 2.5 dpi (Dell’Orso, Juan et al. 2019) (Fig. 2E; and more below), but not in freshly isolated SCs from uninjured muscles (Dell’Orso, Juan et al. 2019) (Fig. S2C). As those prior studies did not focus on *Scx*, its expression might not have been paid attention to. By contrast, our serendipitous finding from ScxGFP mice has led us to document *Scx* expression in activated SCs and regenerative myogenic cells in vivo and in vitro.

**Figure 2:**
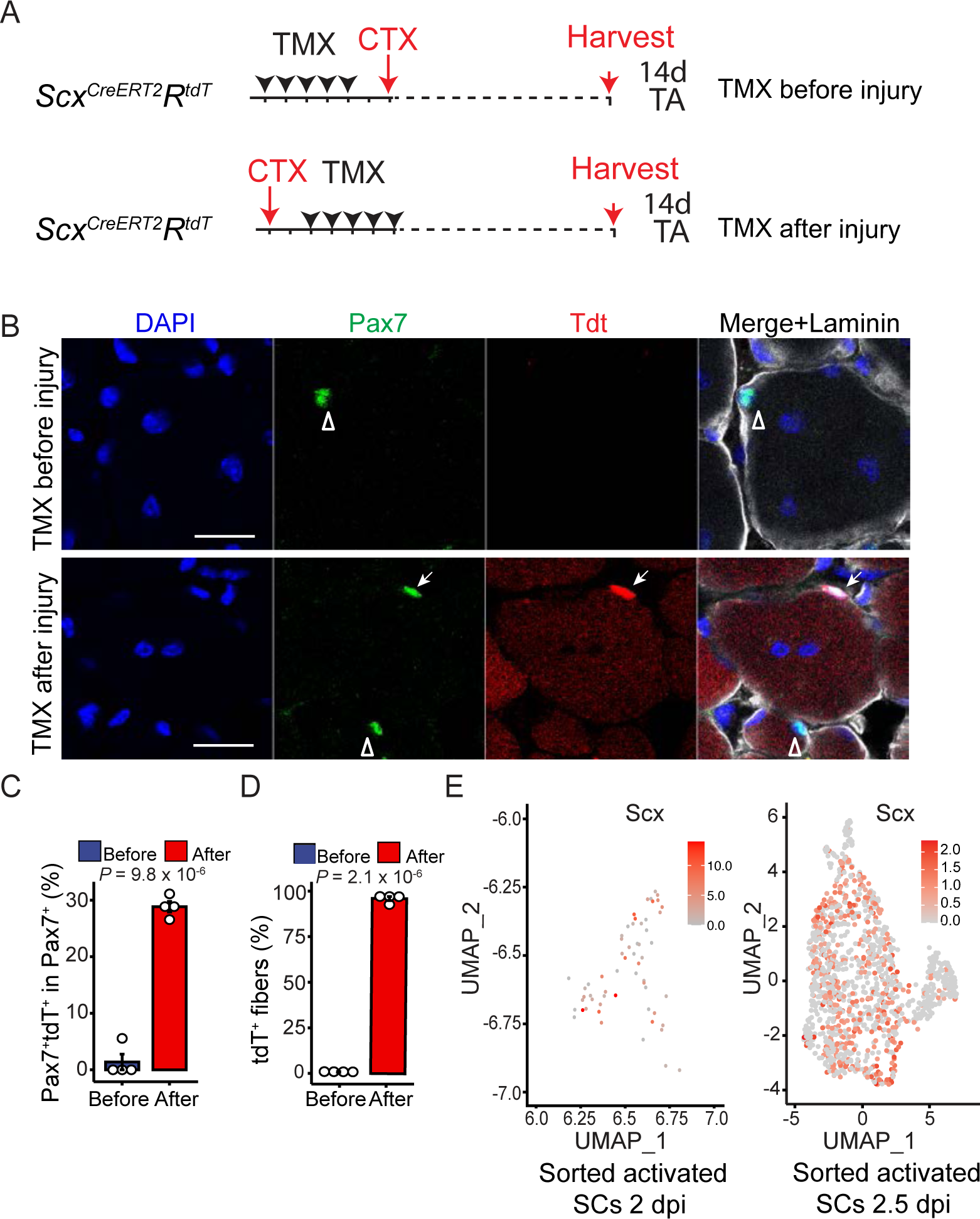
Endogenous *Scx* is expressed by activated but not quiescent SCs. A. Experimental design for *Scx^CreERT2^* mediated inducible lineage tracing with the *R^tdT^* reporter. The two experimental groups are: 1) TMX administrated before injury for 5 days (TMX before injury) and 2) TMX administrated after injury for 5 days (TMX after injury). TA muscles in both groups were harvested at 14d post injury. B. TA muscles from experiment groups in (A) were stained with Pax7 (green) and Laminin (white) and visualized with tdT (no staining). Open arrowheads indicate Pax7^+^ SCs; arrows, Pax7^+^tdT^+^ SCs. C. Percentages of Pax7^+^tdT^+^ SCs in Pax7^+^ SCs examined, from data in B. (N = 4 mice per group; n = 191 (Before, TMX before injury) and 256 (After, TMX after injury) Pax7^+^ cells). D. Percentages of tdT^+^ myofibers in regenerated muscle fibers (with centrally located nuclei), from data in B. (N = 4 mice; n = 1956 (Before, TMX before injury) and n = 2024 (After, TMX after injury) regenerated myofibers). E. Re-analyses for *Scx* expression in two published scRNA-seq data sets of activated myogenic cells at 2 dpi and 2.5 dpi (De Micheli, Laurilliard et al. 2020, Dell’Orso, Juan et al. 2019), displayed by UMAP; colored keys to expression levels are included correspondingly. Nuclei were stained with DAPI; Scale bar = 20 μm. Data are presented with mean ± s.d.; *P* value are indicated. C-D, Unpaired two-tailed Student’s *t*-test were applied.

### *Scx* is required for adult skeletal muscle regeneration

To determine whether *Scx* plays a direct role in the myogenic lineage during regeneration, we combined floxed *Scx (Scx^F^)*(Murchison, Price et al. 2007) and *Pax7^Cre-ERT2^* (*Pax7^CE^*)(Lepper, Conway et al. 2009) to generate ScxcKO mice for TMX-inducible gene inactivation (Fig. S3A); loxP sites flank the first exon of *Scx* (Fig. S3B). Either tdT or YFP (*R^YFP^*) reporter (specified in figures and legends) was included for cell marking. Highly efficient and selective removal of exon1 was determined using genomic DNA samples of FACS-isolated control and ScxcKO SCs (Fig. S3A–D).

### ScxcKO mice have muscle regeneration defects

Next, we injured control and ScxcKO mice with CTX, and compared their regeneration at 5 and 14 dpi (Fig. 3A); samples shown in Fig. 3 carried the tdT reporter. At 5 dpi, ScxcKO’s regenerating myofibers were significantly smaller than those in control mice (Fig. 3B, C). Similar results were obtained in mice carrying the YFP reporter (Fig. S3 J, K). At 14 dpi, regenerated myofibers in ScxcKO mice were still considerably smaller than those of the control (Fig. 3D, E). Thus, *Scx* function is needed in the Pax7^+^ SC-lineage for robust regeneration of muscle fibers.

**Figure 3:**
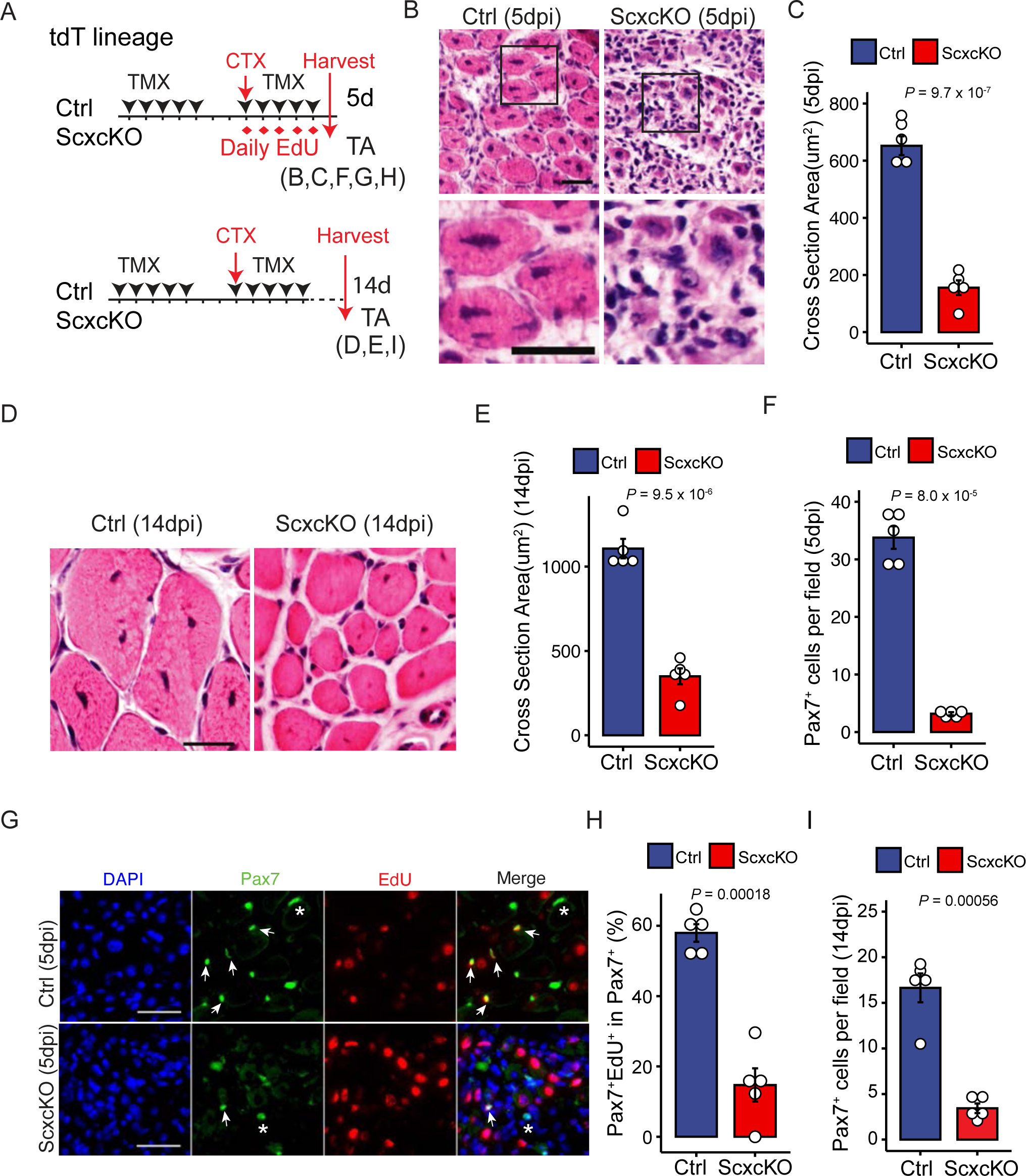
Efficient muscle regeneration requires *Scx* function. A. Experimental designs to compare phenotypes of control (Ctrl) and ScxcKO mice. The *R^tdT^* reporter was included (tdT lineage; see Figure S3A for genotypes). TMX was administered before and after the CTX induced injury to maximize gene inactivation. TA muscles were harvested at 5d or 14d after injury. B-C. (B) Ctrl and ScxcKO TA muscles at 5d post-injury were sectioned and stained with hematoxylin and eosin (H&E) at low (top) and high (bottom) magnifications. (C) Histogram of regenerated muscle fiber cross-sectional area from data in (B). (N = 5 mice per group). D-E. (D) Ctrl and ScxcKO TA muscles at 14d post-injury were sectioned and stained with H&E. (E) Histogram of regenerated muscle fiber cross-sectional area from data in (D). (N = 5 mice per group) F. Histogram of average Pax7^+^ SCs number per imaged field (0.08mm^2^) of TA muscle sections from 5 dpi Ctrl and ScxcKO mice (N = 5 mice per group; n = 2807 Ctrl and n = 442 ScxcKO Pax7^+^ SCs). G. After EdU administration, Ctrl and ScxcKO TA muscles at 5d post-injury were sectioned and stained for Pax7, followed by EdU reaction. Arrows indicate Pax7^+^EdU^+^cells, whereas asterisk, Pax7^+^EdU^-^ cells. Nuclei were stained with DAPI; Scale bar = 20 μm. H. Percentages of EdU^+^ cells within the Pax7^+^ cell population of Ctrl and ScxcKO, from data in G (N = 5 mice per group; n = 858 Ctrl and n = 325 ScxcKO Pax7^+^ SCs). I. Averaged Pax7^+^ SC number per image field (0.4mm^2^) in Ctrl and ScxcKO TA muscle sections from 14d post-injury samples (N = 5 mice per group; n = 350 Ctrl and n = 186 ScxcKO Pax7^+^ SCs). Data are presented with the mean ± s.d.; *P* values are indicated. (C, E-F, H-I) Unpaired two-tailed Student’s *t*-test was applied.

### ScxcKO mice show reduced SC proliferation and renewal

Considering that *Scx* expression is initiated in activated SCs but not in quiescent SCs, and ScxcKO mice have smaller regenerative myofibers, it stands to reason that *Scx* plays a role in their proliferation. At 5 dpi, we noted a ∼ 7-fold reduction in Pax7^+^ SCs in the ScxcKO samples (Fig. 3F, Fig. S3E, F). To show proliferation defect, we administered EdU and assessed cumulative proliferation index over the first 5 days of injury (Fig. 3A, top panel). Compared to the control, the fraction of Pax7^+^ ScxcKO SCs that incorporated EdU (i.e., EdU^+^Pax7^+^) was reduced by ∼ 4-fold (Fig. 3G and 3H). We did not observe appreciable levels of programmed cell death (PCD) in control and ScxcKO at 5 dpi using an anti-cleaved Caspase3 antibody. We also quantified Pax7^+^ SCs number at 14 dpi (Fig. 3A, bottom panel) and found a ∼ 5-fold reduction of renewed SCs in the ScxcKO group (Fig. 3I, Fig. S3H, I). Thus, Scx is autonomously required for SC proliferation and renewal following injury.

When we examined laminin (i.e., basement membrane) and MHC in control 5 dpi samples, we found that the laminin boundary juxtaposed the regenerative myofiber surface (Fig. S3F). As expected, the smaller ScxcKO MHC^+^ fibers did not fill out to the laminin outlines (Fig. S3G). At early injury time points, the laminin pattern represents left-over basement membranes of dead myofibers (caused by injury), i.e., the ghost fiber(Webster, Manor et al. 2016). Ghost fibers are thought to be replaced by basement membranes produced by regenerated fibers over time. At 14 dpi, regenerated myofibers in both control and ScxcKO were tightly surrounded by laminin despite their difference in size (Fig. S3H, I), suggesting that ScxcKO regenerated myofibers are capable of making their own basement membranes.

### Scx is needed for robust proliferation of SC in culture

To examine Scx function in the SC without interactions with other cell types in the injured/regenerative environment, we turned to *in vitro* assays using purified SCs (Fig. 4A). To simplify FACS isolation of SC, we utilized either the tdT or the YFP fluorescent reporter (Fig. S4A, B). SC purity was assessed by staining for Pax7 immediately after FACS. We were surprised that YFP-marked SCs (YFP-SCs) exhibited higher purity than tdT-marked SCs (Fig. 4B). Indeed, Murach and colleagues have reported exosomal transfer of tdT mRNA from lineage-marked Pax7^+^ cells to several other cell types(Murach, Peck et al. 2021). Their finding helps explain lower purity of Pax7^+^ cells by tdT-marking in our hands. We suggest that high levels of tdT mRNA produced by the strong CAG promoter/enhancer leads to more tdT^+^ non-SCs by exosomal transfer, compared to the low-moderate levels of YFP reporter mRNA produced by the Rosa26 promoter.

**Figure 4:**
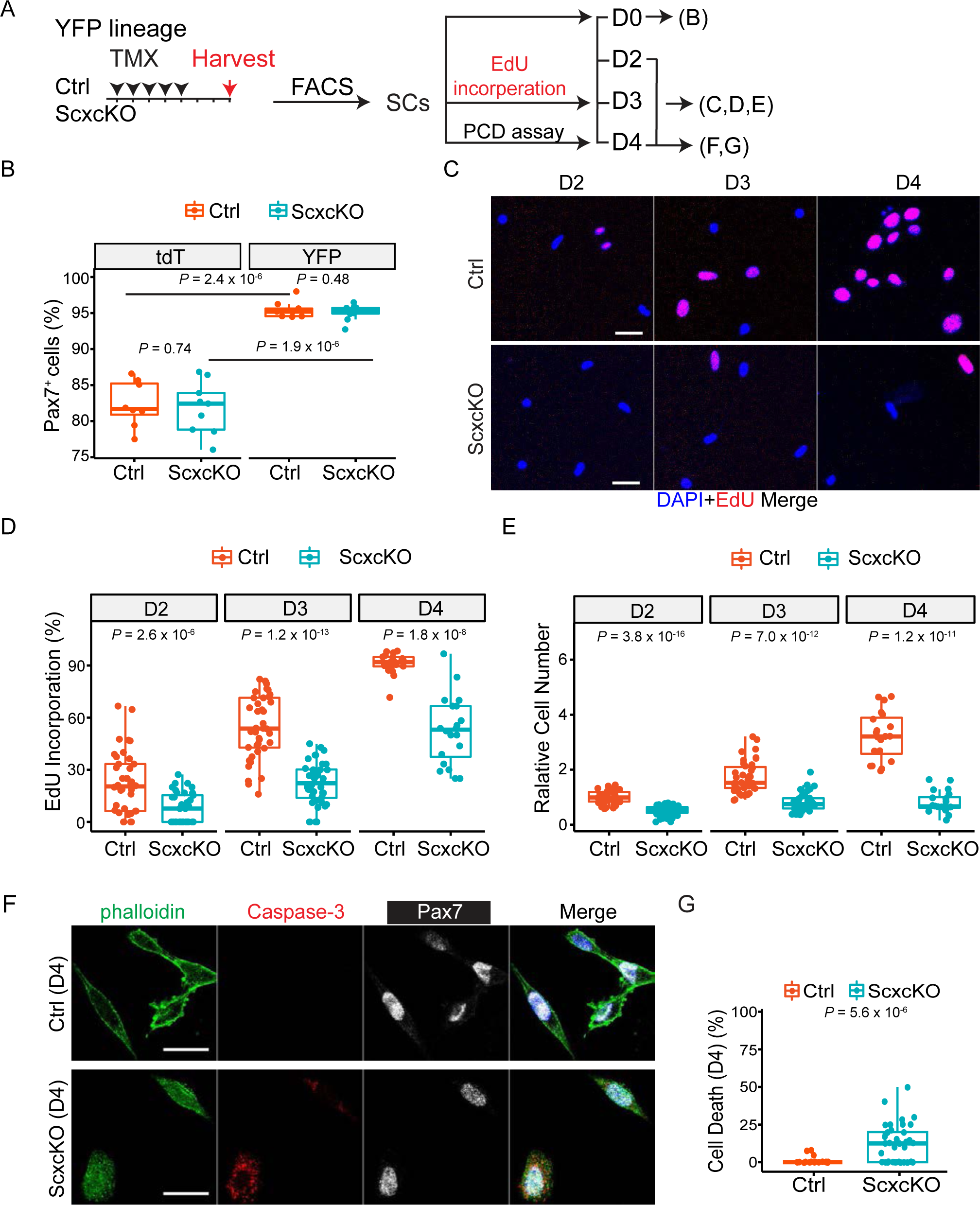
ScxcKO SCs display a proliferation defect. A. Experimental design to obtain YFP lineage marked Pax7^+^ SCs for *in vitro* analyses in (B-F). B. Box plot of percentages of FACS-isolated tdT and YFP marked cells expressing Pax7 by staining immediately after isolation (as D0); each dot represents one image data, 10 image per group, totally n ≥ 1000 cells for each group. C. YFP lineage marked cells were cultured in GM and assayed at days 2, 3, and 4 (D2-D4) intervals. 10 μM EdU was added for 6 h prior to harvesting for EdU detection. D. Box plot of percentages of EdU^+^ cells from data in (C), N=2 mice, each dot represents one image data, 3 well per group, 8 images per well. E. Box plot of ratios of total cell numbers from data in (C); normalized to the average control cell number at D2 as 1. F-G. (F) FACS-isolated Ctrl and ScxcKO SCs were cultured in growth medium for 4 days, harvested, and immuno-stained for Pax7 and cleaved Caspase 3; actin cytoskeleton (to identify cell body) was stained by Phalloidin. (G) Box plot of percentages of cell death (i.e. cleaved Caspase 3^+^ cells) from data in (F); N=2 mice; each dot represents one image data; 2 wells per group, ≥10 images per well; 240 Ctrl and 353 ScxcKO cells examined. Nuclei were stained with DAPI; Scale bar = 20 μm. Data are presented with the mean ± s.d.; adjusted *P* values are shown. (B) Two-way ANOVA; (D-F) Unpaired two-tailed Student’s *t*-test.

As such, we opted to use YFP-marked control and ScxcKO SCs in subsequent studies for higher SC purity. Of note, ScxcKO with YFP reporter had similar regenerative defects as that with tdT reporter (Fig. S3J, K). By EdU incorporation assay, we found that cultured ScxcKO SCs displayed reduced proliferation indices at days 2, 3 and 4. Curiously, the number of ScxcKO cells per imaged area (i.e., cell density) barely increased during this time course, despite EdU incorporation (Fig. 4C, D, E). We next carried out live imaging (Fig. S4C) to document the behavior of control and ScxcKO SCs. Consistent with EdU incorporation, control SCs showed a faster increase in cell number/density than ScxcKO SCs (Fig. S4D, E). ScxcKO cells also showed a slightly reduced cell mobility (Fig. S4F). As we did not observe appreciable levels of PCD using anti-cleaved Caspase3 at 5 dpi, we assessed PCD by another assay (TUNEL) at an earlier time point. Yet, we still failed to detect appreciable TUNEL^+^YFP^+^ myogenic cells in either control or ScxCKO muscles at 3 dpi (Fig. S4I). Intriguingly, we did observe cell loss during live-imaging (Fig. S4G, H): More ScxcKO cells rounded up or appeared necrotic before disappearing. We therefore evaluated PCD of cultured SCs by anti-cleaved Caspase3 and found an increased rate of PCD of ScxcKO SCs, relative to that of the control (Fig. 4F, G, Fig. S4J). Our results support that Scx acts autonomously in the SC to promote proliferation, survival, and migration.

### Scx expression by Single-Cell RNA-Sequencing (scRNA-Seq)

To determine the mechanism underlying Scx’s role in the SC-lineage, we employed scRNA-seq using the 10x Chromium platform (Fig. S5A). For this, multiple sites of BaCl_2_ injection were made to TA and gastrocnemius muscles of control and ScxcKO mice to induce wide-spread injury and activate as many SCs as possible(Morton, Norton et al. 2019). Because the published sc-RNA-seq data(Dell’Orso, Juan et al. 2019) indicated a widespread *Scx* expression at 2.5 dpi (Fig. 2F), we chose this time point for investigation.

YFP-marked control and ScxcKO SCs at 2.5 dpi were FACS-isolated and immediately subjected to scRNA-seq (Fig. 5A). Data were analyzed using the R package Seurat and unsupervised graph-based clustering(Hao, Hao et al. 2021). After filtering, 11,388 control and 12,844 ScxcKO cells, respectively (with ∼23,000 detectable genes), were qualified for analysis. We utilized uniform manifold approximation and projection (UMAP) to display all cells in the unified dataset and performed unsupervised shared nearest neighbor (SNN) clustering to partition cells into 18 (0-17) clusters (Fig. S5B). We annotated the cell types by examining the normalized expression level and frequency of canonical cell type-specific genes. The percentages of cells within each cluster in control and ScxcKO were also calculated (Fig. S5C). Clusters 0-2 and 4-11 contained the majority of cells expressing myogenic genes. Clusters 3 and 12-17 represent non-myogenic cell types, including various immune cells, endothelial cells, and Schwann cells (presumably due to exosomal transfer of YFP mRNA). Cluster 13 was assigned as monocytes/macrophages/platelets, but expressed myogenic genes. They are likely the immunomyoblast proposed by Oprescu et al(Oprescu, Yue et al. 2020). Below, we focused on myogenic clusters to investigate the defects associated with ScxcKO.

**Figure 5:**
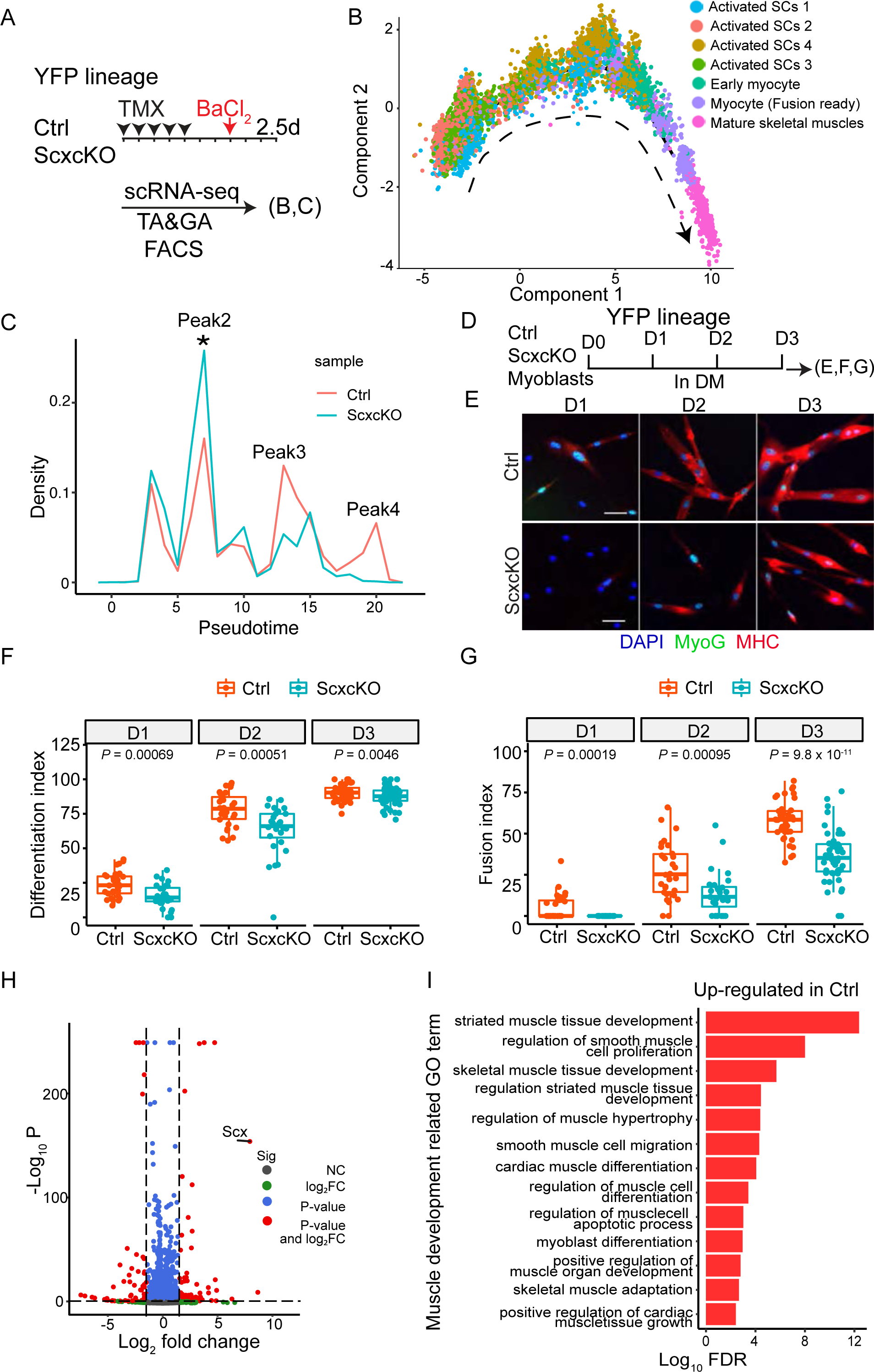
scRNA-seq helps identify the role of *Scx* in myogenic differentiation and fusion. A. SC scRNA-seq scheme for YFP lineage marked SCs. YFP^+^ cells were FACS-isolated from 2.5 dpi BaCl_2_ injured TA and the gastrocnemius (GA) muscles. B. Trajectory analysis of the 7 myogenic clusters (complete cell cluster analysis in Fig. S5) indicated to the right. Arrow indicates the direction of pseudotime trajectory. C. Cell densities of Ctrl and ScxcKO cells along the trajectory in (B). Cell in Peaks 2-4 were used for the DEG analysis; The asterisk indicates Peak 2 as our main focus. D. *In vitro* differentiation assay scheme. SC-derived myoblasts were cultured in GM for 12 h (D0), switched into DM, and harvested daily for analysis over 3 days (D1 - D3). E. Myoblasts subjected the scheme in (D) were stained for MyoG (for differentiation index in F) and for MHC (for fusion index in G). Nuclei were stained with DAPI; Scale bar = 20 μm. F, G. Box plot of differentiation index (F) and fusion index (G) from data in (E). Each dot represents one image data. Unpaired two-tailed Student’s *t*-tests were applied and adjusted *P* values are shown. (N = 3 mice; 3 wells per group per time point; 10 images per well; in total, 3342 control and 2561 ScxcKO cells examined). H. Volcano plot of relative gene expression (Log2 fold change) in Ctrl versus ScxKO cells in Peak 2 (in C). I. GO term enrichment of muscle development related processes from DGEs in (H).

### sc-RNA-seq confirms *Scx* expression during regenerative myogenesis

Of the 11 myogenic cell clusters, we classified them into four categories: early activated SC, activated SC, myocyte, and mature skeletal muscle (Fig. S5B). Within the categories of early activated SC and activated SC, multiple cell clusters were included and numbered as different states. Here, numbers were arbitrarily assigned and not meant to reflect their temporal sequence. Early activated SC 1 - 3 were represented by clusters 4, 5, 9, and expressed varying levels of *Pax7*, *Myod1*, and *Myf5* (Fig. S5B, D, E). We assigned clusters 0, 1, 2 and 7 as activated SC 1 – 4 respectively as they expressed lower levels of *Pax7* (compared to early activated SC). Further evidencing our assignment as activated SCs, more cells in these clusters expressed *Myod1*, *Myf5*, and *Hspa1a*(Francetic T 2011, Senf 2013). Cluster 6 represented early myocyte based on increased expression of *Myog* and *Mef2a*. Cluster 8 cells expressed high levels of *Mymk*, indicating that they are competent for fusion. Cluster 11 cells expressed *Myh1* and *Acta1*, representing mature muscle cell. Cluster 10 cells were unknown myogenic cells for they expressed very low levels of myogenic genes. Among these clusters, the level and cell percentage of *Scx* expression were very low in early activated SCs, and gradually increased from activated SCs to early myocytes, fusion competent myocytes, and mature muscle cells (Fig. S5D, F).

We carried out Monocle 2 trajectory analysis to depict the progression of myogenic cell clusters (Fig. 5B). Given that *Scx* expression is very low in the early activated SC category and we observed ScxGFP only in activated SC experimentally, we excluded early activated SC 1-3 from analysis. The trajectory revealed a time line consistent with our assignment, from activated SC 1 to mature muscle cells. Of the 4 activated SC clusters, activated SC 1 cells were distributed throughout the activation time line up to early myocyte stage, activated SC 2 and SC 3 cells were preferentially located in earlier time line, whereas activated SC 4 cells were found in a later time, revealing their different states. Early myocytes, fusion competent myocytes, and mature muscle cells were ordered as expected.

### scRNA-seq data help identify myogenic differentiation and fusion defects

To understand the timing of *Scx* action, we compared the relative densities of various cell types/states between the control and ScxcKO cells along the pseudotime (Fig. 5C). Relative to control, a higher density of ScxcKO cells, i.e., peak 2 in Fig. 5C, was noted just before their reduction, i.e., peaks 3 and 4. Peaks 3 and 4 correspond to fusion competent myocytes and mature muscle cells, respectively. This information re-directed us to investigate *Scx* function in fusion and differentiation. For this, control and ScxcKO SCs were isolated, cultured, plated at the same density, and then switched to differentiation medium (DM) (Fig. 5D). They were assessed for expression of MyoG (for differentiation index) and MHC (for fusion index) daily over 3 days. More control cells expressed MyoG and MHC when compared to ScxcKO cells at each time point (Fig. 5E, F, G). At day 3, ScxcKO cells caught up in differentiation index (still lower than that of control cells), but were still considerably lower in fusion index. This experimental result, aided by pseudotime analysis, supports a role of *Scx* for regenerative myogenic differentiation.

### Molecular pathways governed by *Scx* in regenerative myogenesis

To gain molecular insight, we examined differentially expressed genes (DEGs) between control and ScxcKO cells along the pseudotime-line. We were particularly intrigued by the DEGs in peak 2, as it represents an early time point of difference to capture candidate direct targets of Scx. There were 3956 DEGs in peak 2 – *Scx* exhibited the largest log_2_ fold change in ScxcKO (Fig. 5H and Table S1). In particular, cyclin-dependent kinases *Cdk1* and *Cdk2* were down-regulated, and CDK-inhibitors *Cdkn1a* and *Cdkn1c* were up-regulated in ScxcKO cells at, and prior to, peak 2 (Fig. S5G). This helps explain the proliferation defects of ScxcKO cells. However, higher cell density with less proliferation potential is somewhat counter-intuitive. We suggest that ScxcKO cells not only proliferate slower but also progress slower towards differentiation and fusion, resulting in their stalling and accumulation at the peak 2 transitional juncture (Fig. 5C).

Consistent with the phenotype of ScxcKO, Gene Ontology (GO) term analysis of peak 2 DEGs revealed that control cells showed enrichment of up-regulated genes in the categories of muscle differentiation, growth, and development, among other pathways overlapping with cardiac muscles (Fig. 5I). 17 genes involved in the muscle cell apoptotic process were found, consistent with increased PCD detected in vitro. Unexpectedly, *Mymk* and *Mymx*, indispensable for myocyte fusion, were expressed higher in ScxcKO than control cells (Tables S1, S2), possibly reflecting their compensatory up-regulation due to compromised differentiation/fusion of *Scx*cKO cells. Analyses of peak 3 and 4 DEGs provide additional information about selective differentiation processes being disrupted in ScxcKO cells (see discussion).

### Identification of direct targets by CUT&RUN assay

To uncover direct gene targets of Scx that regulate muscle differentiation and/or maturation, we utilized the CUT&RUN(Kaya-Okur, Janssens et al. 2020) assay to determine Scx bindings sites in the genome. To aid this endeavor, a triple-Ty1 tag (3XTy1) was fused to the C-terminus of Scx to create a *Scx^Ty1^* allele (Fig. S6A). *Scx^Ty1/Ty1^* mice are viable and fertile without apparent tendon abnormality. Ty1 was detected in linearly arrayed patellar tenocytes (Fig. S6B) and in cultured myoblasts (Fig. S6C, D) derived from *Scx^Ty1/Ty1^* mice. During differentiation time course over 3 days in culture, the largest fraction of cells with detectable Ty1 presented at day 1 (Fig. S6E).

We performed the CUT&RUN using anti-Ty1 on *Scx^Ty1/Ty1^* myoblasts (Scx-CUT&RUN) at 12 h after switching them to DM (Fig. S6F); this time point was chosen to uncover early targets. We included two controls: ScxGFP myoblasts with anti-Ty1, and *Scx^Ty1/Ty1^* myoblasts with non-specific IgG. A total of 1003 binding peaks were identified in 862 gene loci with 33.4%, 38.88% and 22.44% located in intergenic regions, introns, and promoters respectively, alongside other genomic regions, respectively (Fig. 6B; Table S3). These peaks were enriched for the bHLH transcription factor binding motif, the E-box: CAG(A/C)TG (Fig. 6C), indicating high data quality. By integrating the Scx-CUT&RUN data with DEGs in the scRNA-seq data of ScxcKO cells (specifically those in peak 2 of Fig. 5C), we found 207 intersecting genes (Fig. 6D; Table S4). Scx-binding peaks at these gene loci were also enriched for the E-box motif (Fig. 6E), implicating these genes as direct targets. As expected, GO-term of these genes showed enrichment for processes in muscle differentiation, fusion, and myofibril assembly (Fig. 6F; Table S5). We also noted the enrichment for processes of mRNA destabilization, catabolism, poly(A) shortening, etc., suggesting that mRNA metabolism is altered in the ScxcKO (Fig. 6F; Table S5). Four of the 207 candidate direct target genes provide possible explanations for defective differentiation of ScxcKO cells: *Mef2a*, *Capn2*, *Myh9*, and *Cflar* (see Discussion). The Scx-CUT&RUN peaks at these loci were in either the promoter region or intron (Fig. 6G; Fig. S6F), suggesting that Scx binding and/or function is not confined to the promoter region. Taken together, Scx directly regulates a set of E-box containing genes and we discuss how some of these genes help explain the phenotype observed below.

**Figure 6:**
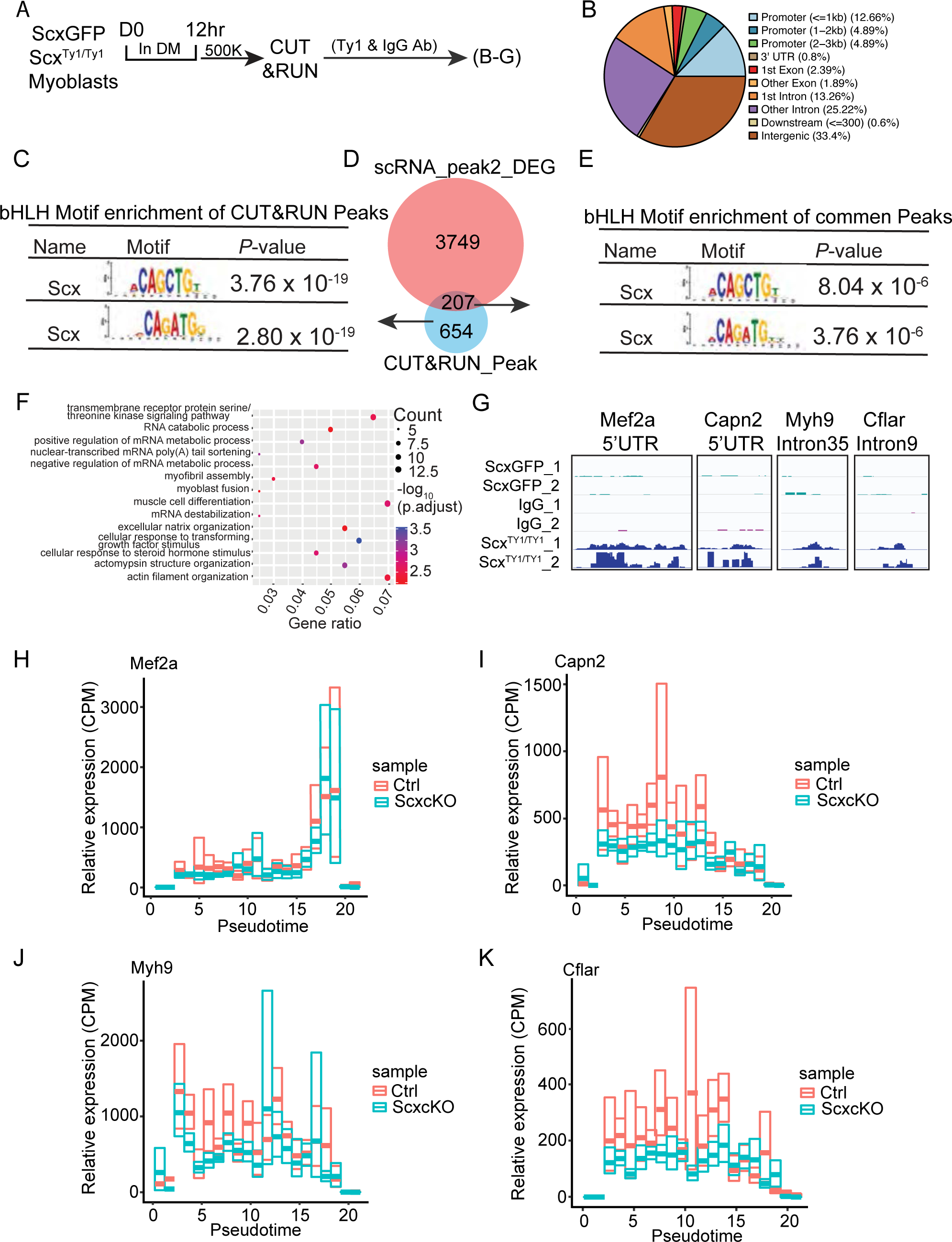
CUT&RUN and scRNA-seq identify direct targets of Scx. A. Experimental scheme for CUT&RUN profiling of the Scx binding in the genome of Scx^Ty1/Ty1^ and ScxGFP myoblasts. Primary myoblasts derived from SCs of Scx^Ty1/Ty1^ (experimental group) and ScxGFP (control group) mice were used. They were cultured in GM for 12 h (D0), and switched to DM for 12 h for use. 500,000 (500K) cells per group were subjected to CUT&RUN using an anti-Ty1 antibody or an IgG control antibody, in duplicate. B. Pie chart for distribution of Scx CUT&RUN peaks in various regions of the genome. C. Motif enrichment analysis with SEA from MEME suite (v. 5.5.0) identified bHLH protein binding motif (i.e. E-box) in all Scx CUT&RUN binding peaks. D. Venn diagram of intersecting genes (207 genes) between Scx CUT&RUN target genes (861) and DEGs (3749) in Peak 2 of Fig. 5C. E. Motif enrichment analysis (as in C) of the 207 genes in (D) also showed enrichment of bHLH protein binding motifs, the E-box. F. GO term analysis of the 207 genes in (D). GO terms with *P* < 0.0001 were plotted. G. Genomic snapshots of Scx CUT&RUN peaks on 4 select genes related to muscle differentiation. H. Expression levels (CPM, counts per million UMI) of the 4 select genes in (G) along the pseudotime trajectory (same trajectory as Fig. 5C).

## Discussion

Here we show that *Scx* is expressed in activated mouse SCs, and it regulates many aspects of muscle regenerative process, from proliferation, cell survival, migration, to differentiation and fusion. The Scx target genes we identified underscore its function in muscle regeneration.

### The multiplicity of Scx-lineage

Since the initial description of the *Scx* gene(Cserjesi 1995), most efforts have been focused on its role in tendon. Its early expression in the syndetome and the limb mesenchyme eventually becomes realized in tendons, ligaments, and CT(Schweitzer, Chyung et al. 2001, Brent, Schweitzer et al. 2003, Tozer and Duprez 2005, Pryce, Brent et al. 2007). Lineage tracing by *Scx^Cre^* confirmed aforementioned descendant cell types alongside other cell types(Esteves de Lima, Blavet et al. 2021, Ono, Schlesinger et al. 2023). Of relevance, a lineage contribution to myofibers was found. The temporal emergence of Scx^+^ cells with myogenic potential was not provided by constitutive Cre-mediated lineage tracing. On the other hand, TMX-inducible lineage tracing mediated by the CT marker gene *Ors1* (i.e., using an *Ors1^CreERT2^*) revealed myogenic incorporation competence that declines towards late embryogenesis(Esteves de Lima, Blavet et al. 2021). A Prx1^+^ CT population has also been shown to incorporate into the myofiber near the myotendonous junction (MTJ) at neonatal stages(Yaseen, Kraft-Sheleg et al. 2021). Consistently, scRNA-seq of embryonic chick limb mesenchyme identified a cell cluster co-expressing CT and myogenic signatures at multiple stages(Esteves de Lima, Blavet et al. 2021). Whether these bi-potential CT/myogenic cells arise from dermomotyotome, syndetome or a yet-to-be identified origin, remains to be rigorously examined.

We show here that adult SCs express ScxGFP upon injury and culture, and that ScxGFP is co-localized with Pax7, MyoD and MHC. scRNA-seq data confirm endogenous *Scx* expression in multiple regenerative myogenic clusters/states, in which the other CT markers *Twist2*, *Ors1*, and *Pdgfra* are barely detectable. Moreover, only the lineage-marked Scx^+^ cells induced after, but not prior to, injury contribute to regenerative muscles and SCs. Together, these results support that muscle interstitial Scx^+^ CT (lineage-marked prior to injury) have no myogenic potential, whereas activated Pax7^+^ SCs expressing *Scx* (lineage-marked after injury) can contribute to new muscles and SCs. This is consistent with transplanted Scx^+^ CT(Giordani, He et al. 2019) lacking a contribution to muscle. Adult muscle interstitial CT are highly heterogeneous within a muscle group as well as between muscle groups based on scRNA-seq data, and not all CT express *Scx*(Muhl, Genove et al. 2020). Anatomically, adult muscle interstitial *Scx*^+^ cells are paramysial cells that line the perimysium(Muhl, Genove et al. 2020). Lineage tracing data showed that Scx^+^ CT and MTJ cells were descendants of Hic1^+^ MPs, but no myofiber incorporation from the Hic1^+^ lineage was noted(Scott, Arostegui et al. 2019). Whether CT/myogenic bipotential progenitors exist in adult muscle is of considerable interest. Regardless, our results strongly support that SCs express *Scx* after activation and require *Scx* function for efficient regeneration.

### *Scx* function in tendon versus muscle

*Scx* has been consider a master regulator of tendon (and ligament) development as *Scx* mutant mice develop severely compromised tendons in the limbs and tail(Murchison, Price et al. 2007, Yoshimoto, Takimoto et al. 2017, Shukunami, Takimoto et al. 2018). *Scx* is required for the expression of multiple tendon matrix protein encoding genes such as *Col1a1*, *Col3a1*, and *Tnmd* (Shukunami, Takimoto et al. 2018), but not for tendon progenitor specification. Ablation of embryonic Scx^+^ cells led to mis-patterned muscle bundles(Ono, Schlesinger et al. 2023), supporting an interdependence between muscle and tendon for connectivity(Kardon 1998). Retrospectively, the observed muscle mis-pattern by ablating Scx^+^ cells likely included ablation of CT/myogenic cells and tendon cells. In adult, *Scx* continues to be required for tendon growth and repair after injury(Howell, Chien et al. 2017, Sakabe, Sakai et al. 2018, Gumucio, Schonk et al. 2020, Korcari, Muscat et al. 2022). By contrast, we focused on *Scx* function in proliferation, migration, differentiation, and fusion within the Pax7^+^ SC lineage for muscle regeneration.

### Downstream genes with implication to the myogenic defect of ScxcKO

GO-term analyses and literature reviews of the 207 DEGs from our scRNA-seq and CUT&RUN data sets identified genes in myogenic processes, instead of genes in tenogenic or CT processes. Several of these downstream genes help us understand how *Scx* may act to regulate regenerative myogenesis: *Mef2a*, *Capn2*, *Myh9*, and *Cflar*. Knocking-out and knocking-down *Mef2a* led to compromised myoblast differentiation *in vivo* and *in vitro*, respectively (Seok, Tatsuguchi et al. 2011, Liu, Nelson et al. 2014, Estrella, Desjardins et al. 2015, Wang, Yang et al. 2018). Reduced *Mef2a* levels explain the compromised myoblast differentiation of ScxcKO cells. Consistently, several *Mef2a* target genes, such as *Hspb7*, *Atp1a2*, *Tmem182*(Wales, Hashemi et al. 2014) were also down regulated (Table S1, S6). Capn2 is a calpain isoform expressed in the skeletal muscle, and the locus harbors 5 E-boxes and 1 MEF-2 binding sites(Dedieu, Mazeres et al. 2003). Knocking-down *Capn2* in C2C12 cells led to compromised cell migration and fusion(Honda, Masui et al. 2008), as observed for ScxcKO cells. *Myh9* was shown to regulate bi-polar cell morphology and alignment during myocyte fusion *in vitro*(Swailes, Colegrave et al. 2006). Its down regulation is consistent with the defective fusion of ScxcKO cells. Lastly, *Cflar* were shown to proliferation and prevent apoptosis in vascular smooth muscle cells and T lymphocytes(Wang, Prince et al. 2002, Zhang and He 2005, Budd, Yeh et al. 2006, Vesely, Heilig et al. 2009). It may act similarly in the SC to explain reduced proliferation and increased cell loss of ScxcKO cells. These 4 genes displayed reduced expression levels at the early part of the pseudotime trajectory (Fig. 6H-K), consistent with them being direct targets. The other 203 genes likely also contribute to aspects of the ScxcKO phenotype in ways yet to be determined. Taken together, Scx directly regulates a set of E-box containing genes, and several of these genes have direct implications to the phenotype observed.

### Scx downstream target genes in tendon versus muscle

As bHLH proteins, both Scx and MyoD1 bind E-box, CANNTG; the central two nucleotides distinguish binding affinities for different bHLH proteins. The initial characterization of Scx showed that it only binds to the left E-box (CATGTG) in the enhancer (with 10 E-boxes) of the muscle creatine kinase (*MCK*) gene(Cserjesi 1995), whereas MyoD1 has higher affinity to the right E-box (CACCTG). Not surprisingly, the right E-box (with high affinity for MyoD1) is more important than the left E-box for *MCK* expression (Nguyen, Buskin et al. 2003) By contrast, characterization of the promoter of a tendon-specific gene *Tnmd* identified two Scx-responsive E-boxes, CAGATG and CATCTG (Shukunami, Takimoto et al. 2006, Shukunami, Takimoto et al. 2018). Our CUT&RUN identified CAG(A/C)TG as high-ranking Scx binding motifs in myogenic cells, which is the same as one of the E-boxes in the *Tnmd* promoter. Recently, bulk-RNA-seq and ChIP-seq were combined to define *Scx* target genes in embryonic tenocytes(Li, Wu et al. 2021). Although their and our data sets are not age-matched and obtained by different methods, we compared them nonetheless. Overall DEGs (including those without Scx-binding sites) between our and their data yielded minimal overlap (0.9%, using the criteria of log2FC > 0.5). Two genes, *Htra3* and *Olfml2b*, are overlapping DEGs (48 genes for tenocytes and 207 genes for myoblasts) with Scx-binding sites, and neither gene has been studied in tendon or skeletal muscle. Importantly, the compiled E-box sequences bound by Scx in tenocytes and myoblasts are not different, i.e., CAG(A/C)TG. Thus, the deployment of *Scx* by adult SCs is not a re-use of its function in the tendon. The distinctiveness of Scx target genes between these two tissues is most likely attributed to chromatin accessibility imposed by different epigenomes.

### Indirect target genes of Scx further explain defects of the ScxcKO

Although we emphasized Scx’s direct target genes in peak 2 of Fig. 5C, there were many more DEGs that were indirect targets (i.e. without significant Scx-CUT&RUN peaks). Dysregulation of those genes also provide insights to *Scx*-regulated muscle regeneration. For the proliferation defect, we mentioned 4 dysregulated cell cycle regulators in the results section. In addition, *Erk1/2/3* (*Mapk1/3/6*), known for their role in cell growth, also exhibited lower expression levels in ScxCKO cells during the early pseudotime phase (Tables S1, S6). For GO-enrichment in cell migration (20 genes; Table S2), *Itga2* and *Crk* are worth noting as they have been shown to play this role in non-muscle cell contexts(Ren, Zhao et al. 2019, Cai, Guo et al. 2022), (Chuang, Wu et al. 2018), (Huang, Clarke et al. 2015). They may mediate myogenic cell migration under the umbrella program of *Scx*. For differentiation and fusion at the later pseudotime phase, there are 1942 and 755 DEGs (Tables S6, S7) in peaks 3 and 4 (Fig. 5C), respectively. GO-term analyses identified 74 (in peak 3) and 47 (in peak 4) genes related to muscle differentiation (Tables S8, S9). Several of them have documented roles in myogenic differentiation, e.g., *Hacd1*(Lin, Yang et al. 2012), (Blondelle, Ohno et al. 2015)*, Klhl41*(Paxton, Cosgrove et al. 2011, Ramirez-Martinez, Cenik et al. 2017)*, Ehd2*(Doherty, Demonbreun et al. 2008, Posey, Pytel et al. 2011), and *Lmna*(Dubinska-Magiera, Zaremba-Czogalla et al. 2013, Maggi, Carboni et al. 2016). As these gene products act in different cellular compartments and mediate distinct processes, *Scx* does not appear to govern a singular process for muscle differentiation. How these indirect genes come to be dysregulated in the absence of *Scx* remains to be deciphered.

Together with the embryonic CT/myogenic bipotential cells and the Prx1^+^ CT capable of myogenic fusion near the MTJ, our results add an additional layer of complexity and further blur the molecular and cellular boundaries that divide muscle versus tendon/CT identity. The wealth of information of heterogeneous cell types and states obtained by scRNA-seq will continue to break many long-accepted concepts of tissue-restricted functions of transcription factors.

## Methods

### Mouse strains

*Pax7^CE/+^* (*Pax7^Cre-ERT2^*)*(Lepper, Conway et al. 2009)*, *R^YFP^* (Gt(ROSA)26^Sortm19EYFP)Cos/J^)^(S Srinivas 1 2001)^, *R^tdT^* (Gt(ROSA)26^Sortm14(CAG-tdtomato)Hze/J^)^(Madisen, Zwingman et al. 2010)^, *Scx^F^* (*Scx^tm1Stzr^*)*^(Murchison, Price et al. 2007)^*, *Scx^CreERT2^* (*Scx^tm2(cre/ERT2)Stzr^*)*(Howell, Chien et al. 2017)* and Tg-ScxGFP(Pryce, Brent et al. 2007) alleles were obtained from either original investigators or the Jackson Laboratory (JAX). *Scx^Ty1^* allele was made and characterized by our group, with 3 Ty1 tags (EVHTNQDPLD) inserted upstream of the TGA codon of the *Scx* gene. All animals had mixed background. Genotypes of animals are stipulated in text, figures and legends. For qPCR to determine ScxcKO efficiency, primers are in Table S10 (referenced in Fig. S3 legend). Both sexes were used in all experiments and grouped together, except that, only males were used for scRNA–seq. All mice were used between 2-4 month of age. All animal treatment and experiments were approved by the Institutional Animal Care and Use Committee (IACUC) of the Carnegie Institution of Washington (Permit number A3861-01).

### TMX and EdU administration

Tamoxifen (TMX; Sigma) was prepared as 20 mg ml^−1^ stock in corn oil (Sigma) and administered by intraperitoneal injection at the mice at 4mg per 40g body weight following regimens in text, figures and legends. For daily *in vivo* proliferation tracing, 5-ethynyl-2′-deoxyuridine (EdU, 0.5 mg/ml in PBS; Thermo Fisher Scientific) was administered by intraperitoneal injection at 0.1 mg per 20 g body weight per injection. Muscle samples were collected as specified in figures and legends.

### Muscle injury

For CTX injury, control and experimental mice were anaesthetized by isoflurane/oxygen vapor, tibialis anterior (TA) muscle was injected with 50 μl of 10 μM CTX (Cardiotoxin, Sigma-Aldrich) using an insulin syringe (U-100; BD); For BaCl_2_ injury, control and experimental mice were anaesthetized with 2,2,2-tribromoethanol (Sigma) which was prepared as a 100% (w/v) stock solution in 2-methyl-2-butanol (Sigma), diluted 1:40 in PBS, This anesthetic was delivered through intraperitoneal injection at 10 μl per 1 g body weight. Muscle injury was administrated by injecting 2-4 ul per site of 1.2% (w/v) barium chloride (Fisher Chemical) into approximately 25 sites in the lower hindlimb muscles. Animals were then harvested at the post-injury time point stated in the text and figure legend.

### Muscle sample preparation

TA muscle samples were collected, fixed for 8 min in ice-cold 4% paraformaldehyde (PFA) (EM Grade, cat, 157-4) in PBS, sequentially incubated in 10, 20 and 30% sucrose/PBS overnight, embedded in OCT compound (Tissue-Tek, #4583), frozen in isopentane (Sigma)/liquid nitrogen and stored at −80 °C until cryosectioning. Cross-sections (10μM) of the mid-belly region of the muscle were stained with haematoxylin and eosin (H&E; Surgipath) or used for immunostaining and EdU reactions.

### SC isolation by FACS and myoblast culture

SCs were isolated according to the protocol described previously(Liu, Cheung et al. 2015, Yue, Wan et al. 2020) with slight modifications. Briefly, mouse hindlimb muscles were dissected, minced and digested with collagenase II (1000U/ml, Worthington) in wash medium (10% Horse Serum (HS, Invitrogen)) in Ham’s F-10 medium with 1% penicillin/streptomycin (P/S, Gibco)) for 1.5h followed by centrifugation and washing. Then, the tissue slurry was further digested by collagenase II (100U/ml) and dispase (1.1U/ml, Gibco) in wash medium for 0.5 h to get single cell suspension for cell sorting. The cell suspension was sorted using a BD ARIA III sorter equipped with 375-nm, 488-nm, 561-nm, and 633-nm lasers. For Fluorescence sorting the YFP^+^ (tdT^+^) cells are sorted with green fluorescence; FITC channel 488-nm (red fluorescence; PE channel 568-nm). For 4 surface makers labelling, cells were incubated with 4,6-diamidino-2-phenylindole (DAPI) and fluorophore-conjugated antibodies (BioLegend) against CD31, CD45, stem cells antigen-1(Sca1) and vascular cell adhesion protein 1 (Vcam1) at 4 °C for 0.5 h. After washing, cells were subjected to FACS (DAPI^-^, CD31^-^, CD45^-^, Sca1^-^ and Vcam1^+^ cells were collected), and data were collected by FACS Diva software v.6.1.3 (BD Biosciences). A small fraction of sorted cells was immunofluorescence staining for the muscle stem cell markers Pax7. For short-time cell culture, freshly sorted mononuclear SCs were plated on Matrigel (catalogue no. 354248; Corning) coated dishes (37 °C for 1 h) and cultured in SCs culture medium (growth medium: 20% FBS, 5% horse serum, 1% penicillin/streptomycin, 1% GlutaMAX supplement (Gibco) and 2.5 ng/ml FGF (R&D systems) in DMEM (Gibco)) at 37°C in tissue culture incubators with 5% CO_2_. Cells were harvested as specified in the text and figure legend; For long-time cell culture to get stable primary myoblast cell line, freshly-isolated satellite cells were cultured in Ham’s F10 (F10, Sigma), 10% HS and 1% P/S for 2 days, then passage the cell into culture medium and expended the cell for 2 more passages. After that the cells were Cryopreserved into liquid nitrogen for later CUT&RUN and myoblast differentiation and fusion assays; For *in vitro* differentiation and fusion assay, freshly sorted cells or frozen cells were thawed and cultured in growth medium for 12 h then changed into differentiation medium (2% HS, 1% P/S in DMEM) on Matrigel coated plate and harvested as specified in the figure legend; for EdU labelling, 10μM EdU was added to the SC culture medium for 6hr before harvesting for assay.

### Live imaging

Freshly isolated SCs were cultured in growth medium on Matrigel-coated 48well dish at 5K cells per well, 3 wells per sample, and 5 locations per well. Images were collected every 10 min for 4 days. A short interval at the end of each day was used to adjust the focus and add medium to get quality video and keep the cell in a good state. The videos were collected with a Nikon Ti2 system.

### Immunofluorescence staining and detection

Muscle sections were hydrated with PBS, permeabilized with 0.5% Triton X-100 (Sigma-Aldrich)/PBS (0.5% PBT) for 15 min, washed with 0.05% PBT, and blocked with MOM block (Vector Lab) overnight. Sections were washed and incubated in blocking solution (1 X carbo-free blocking solution (Vector Lab)) and 10% goat serum in 0.05% PBT) for 2 h at room temperature, followed by incubation with primary antibodies diluted in blocking solution overnight at 4°C. Sources and dilution for primary antibodies are provided in Table S11, Sections were then washed with 0.05% PBT 3 times and incubated with appropriate Alexa Fluor-conjugated secondary antibodies (1:1,000 for Alexa 488 and Alexa 568 and 1:500 for Alexa 647; Thermo Fisher) in blocking buffer for 1 h at room temperature. Sections were then washed with 0.05% PBT, stained with DAPI (1μg/ml in 0.05% PBT) and mounted in anti-fade diamond solution (Invitrogen). tdT fluorescence was preserved, so no antibody staining was used. This protocol was also used for SCs and myoblasts with two modifications, 1) the cells were fixed for 10 min in 4% PFA and 2) cells were only blocked with blocking buffer (10% goat serum in 0.05% PBT). For EdU detection, the Click-iT Reaction Kit (Thermo Fisher Scientific) was used before blocking according to the manufacturer’s recommendations.

### TUNEL assay

The TUNEL assay kit (Cell Signaling #48513; Fluorescence, 594) was procured from Cell Signaling Technology, and the assay was performed according to the provided protocol with minor modifications. Briefly, pre-fixed frozen sections were rinsed and permeabilized as described in the immunofluorescence staining protocol. Subsequently, each section was incubated with TUNEL Equilibration Buffer for 5 minutes. Following the removal of the Equilibration Buffer, sections were immediately incubated in 50 µL of TUNEL reaction mix, prepared by adding 1 µL of TdT Enzyme to 50 µL of TUNEL Reaction Buffer, for 2.5 hours at 37°C. The sections were then rinsed three times in PBST for 5 minutes each. Samples were either mounted or further processed for immunostaining with immunofluorescence staining protocol.

### CUT&RUN

CUT&RUN experiments were carried out according to the CUTANATM CUT&RUN Protocol vison1.8 with modifications. Briefly, 500k cells were used for each sample and 0.01% digitonin (w/v) was used during the whole process. The antibodies used in the procedure were provided in Table S11. For the library preparation and sequencing, ThruPLEX DNA-seq Kit (Takara) was used to construct the CUT&RUN DNA library for sequencing on an Illumina platform. 5-10 ng purified CUT&RUN-enriched DNA was used for the library preparation. The whole process was performed according the protocol with deviations aiming to preserve short DNA fragments (30-80 bp). After the Library Synthesis step (adaptor ligation), 1.8 x volume of AMPure XP beads was added to the reaction to ensure high recovery efficiency of short fragments. 12 cycles of PCR amplification system was used, then the reaction was cleaned up with 1.2 x volume of AMPure XP beads. The libraries were assayed with a High Sensitivity DNA bioanalyzer (Agilent) for quality control and sequenced in the Illumina NextSeq500. To enable determination of fragment length, paired-end sequencing was performed (2x75 bp, 8 bp index). The data is analyzed by nf-core/CUT&RUN pipeline with –seacr stringent parameters (version 2.0)(Ewels, Peltzer et al. 2020). The overlapped peaks between replicates were considered as conserved peaks and used for downstream analysis. The motif analysis is carried out with SEA from MEME suit (v. 5.5.0) to identify the enrichment of bHLH family binding motifs in CUT and RUN targets.

### Microscopy and image processing

H&E staining images of TA muscle sections were captured by a Nikon 800 microscope with X20 Plan Apo objectives and with a Canon EOS T3 camera using EOS Utility image acquisition software v.2.10. Fluorescent images of TA muscle sections and cultured myoblasts were either captured by a Nikon Eclipse E800 microscope equipped with X20/0.50 Plan Fluor, X40/0.75 Plan Fluor and Hamamatsu C11440 digital camera using the Meta Morph Microscopy Automation and Image Analysis Software v.7.8.10.0, or captured by a Leica SP5 confocal microscope equipped with a X63/1.4 Plan Apo oil objective using the Leica Application Suite Advanced Fluorescence software version 2.7.3.9723. The same exposure time was used and the images were processed and scored in blinded fashion using ImageJ v.64 (National Institutes of Health (NIH)). If necessary, brightness and contrast were adjusted for an entire experimental image set. Cell number, fiber diameter, fiber number and fiber cross-sectional area were measured with ImageJ v.64.

### Single cell RNA sequencing (scRNA-seq)

The lower hindlimb muscles (TA and gastrocnemius muscles) of the ScxcKO and control mice were injured with BaCl_2_ and recovered for 2.5 day. Pax7 lineage cells were FACS-isolated by YFP fluorescence. Cells were suspended in PBS and counted by hemocytometer into 1000 cells/μl. Around 17,000 cells per sample were used for single-cell library preparation using the 10x Genomics platform with Chromium Next GEM Single Cell 3′ GEM, Library and Gel Bead Kit v.3.1 (PN-1000121, v.3 chemistry), Single Cell 3′ A Chip Kit (PN-1000009) or Chromium Next GEM Chip G Single Cell Kit (PN-1000127), and i7 Multiplex Kit (PN-120262). We followed the 10x protocol exactly to prepare the scRNA-seq library. In brief, for v.3 chemistry, 16.5μl cell suspension and 26.7μl nuclease-free water were mixed with 31.8μl reverse transcription master mix. Of this 75μl mix, 70μl was loaded into the Chromium Next GEM Chip G. After barcoding, cDNA was purified and amplified with 11 PCR cycles. The amplified cDNA was further purified and subjected to fragmentation, end repair, A-tailing, adaptor ligation and 14 cycles of sample index PCR. Libraries were sequenced using Illumina NextSeq 500 for paired-end reads.

### Analyses of scRNA-seq data

Sequencing reads were processed with the Cell Ranger version 6.0.1 (10X Genomics, Pleasanton, CA) using the mouse reference transcriptome mm10. From the gene expression matrix, the downstream analysis was carried out with R version 4.0.2 (2020-06-22). Quality control, filtering, data clustering and visualization, and the differential expression analysis were carried out using Seurat version 4.0.3 R package(Hao, Hao et al. 2021). Cells with <1000 UMIs or mitochondrial reads >10% were removed from the analysis. In addition, we removed potential doublets by DoubletFinder (v. 2.0.3)(McGinnis, Murrow et al. 2019). After log-normalizing the data, the expression of each gene was scaled regressing out the number of UMI and the percentage of mitochondrial genes expressed in each cell. The two datasets were integrated with IntegrateData function from Seurat. We performed PCA on the gene expression matrix and used the first 20 principal components for clustering and visualization. Unsupervised shared nearest neighbor (SNN) clustering was performed with a resolution of 0.6 and visualization was done using uniform manifold approximation and projection (UMAP). The Scx expressed myogenic lineage clusters 0, 1, 2, 6, 7, 8, and 11 were subjected to trajectory analysis by monocle 2 (v. 2.16.0)(Qiu, Mao et al. 2017). To organize cells in pseudotime, we performed new dimension reduction and regressed out mitochondrial effects with reduceDimension function and unsupervised cluster them into 5 clusters with clusterCells function. The differentially expressed genes were calculated by differentialGeneTest and the top 500 differentially expressed genes are used to order then used by Monocle for clustering and ordering cells using the DDRTree method and reverse graph embedding.

## Data availability

Mouse single-cell RNA sequencing data and CUT&RUN data were uploaded to NCBI (PRJNA1050758). Select intermediate RDS objects are available at figshare (https://figshare.com/projects/Skeletal_Muscle_Satellite_Cells_Co-Opt_the_Tenogenic_Gene_Scleraxis_to_Instruct_Regeneration/190935)

## Quantification and statistical analysis

Statistical analyses were performed in R version 4.0, with tidyverse and ggplot2 packages. The statistical significance of results was determined by unpaired Student’s t-test and Two-way ANOVA.

## Acknowledgement

We thank Eugenia Dikovsky and mouse facility crew for animal housekeeping, Allison Pinder for technical assistance in scRNA-seq, Mahmud Sidiqqi for assistance in microscopy, and L. Yue for assistance with FACS of SCs. This work is supported by the NIH (AR060042 and AR071976) and Carnegie fund to CMF.

## Author contributions

T. H and C.-M. F. made the initial observation, collected critical preliminary data, and conceptualized the projects. Y.B. and C.-M.F. designed experiments, analyzed data, made conclusions, and wrote the manuscript. C.B. helped in genotyping the mouse strains, data collection and quantification. Y.B. and M.H. performed bioinformatic analyses. All authors contributed to writing and editing.

## Competing interests

The authors declare no competing interests.

## Data and/or Code Availability

All data that support the findings of this study and the custom code used in this study are available from the corresponding authors upon reasonable request.

---

**Figure S1:**
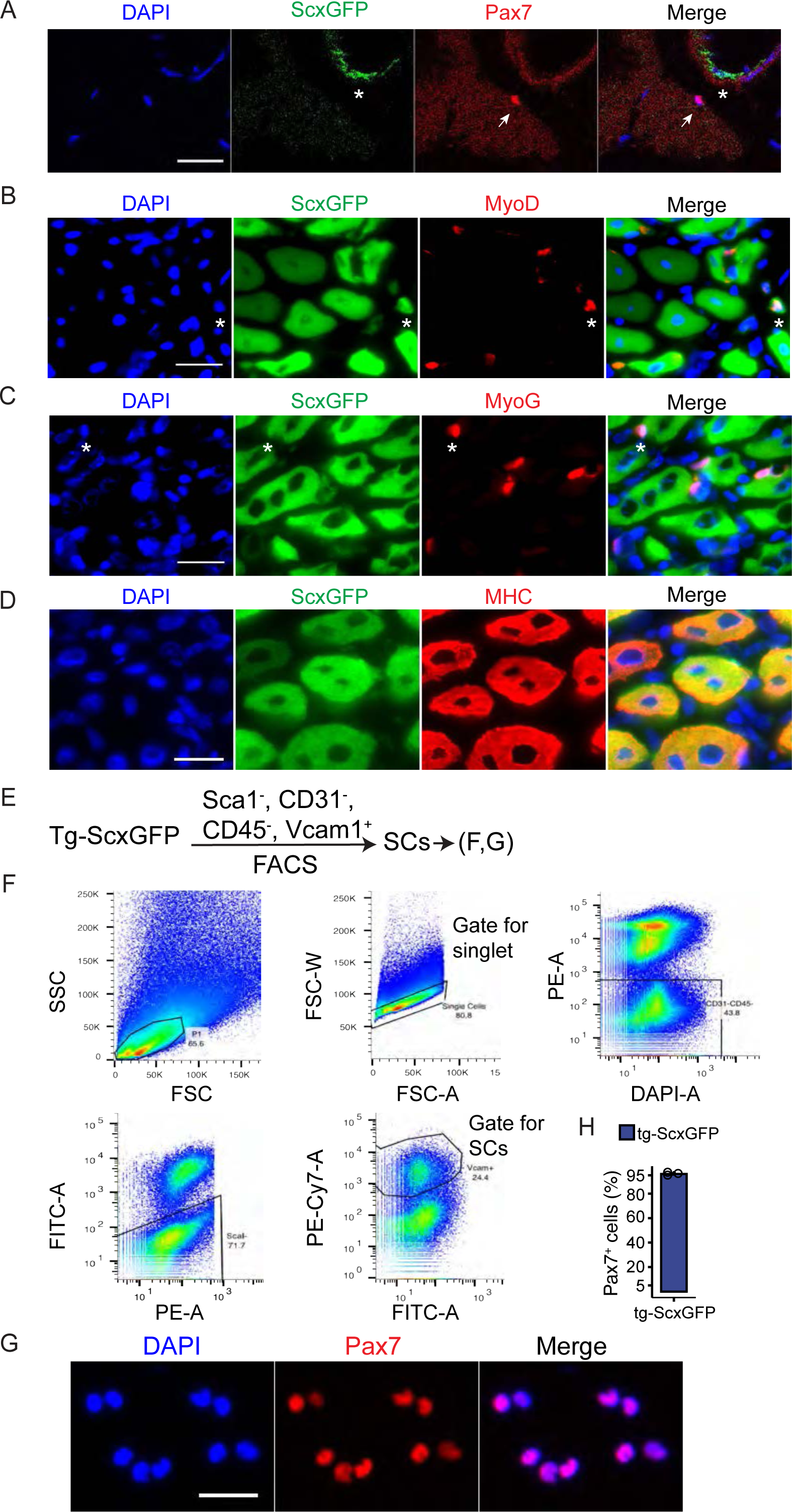
Addition data for Figure 1. A. TA muscle from uninjured ScxGFP mice were sectioned and stained with Pax7 and GFP (N = 4 mice; n = 203 Pax7^+^ cells examined and no Pax7^+^GFP^+^ cells found). The arrow indicates the Pax7^+^ SC; asterisk, ScxGFP^+^ intramuscular CT. B-D. Split channel images of data in Figure 1C; asterisks indicate the same cells. E-F. FACS strategy (E) and profiles (F) to support Fig. 1E. (F) FACS plot and population hierarchy of SC from 4 surface markers sorting in pseudocolor plots. G. Freshly isolated SCs from ScxGFP mice were cyto-spun and stained with Pax7 and DAPI. H. Percentage of Pax7^+^ cells in freshly isolated ScxGFP SCs by FACS procedures in (E, F). (N = 3 mice; n =1805 cells). Scale bar = 20 μm. Data are presented with mean ± s.d.

**Figure S2:**
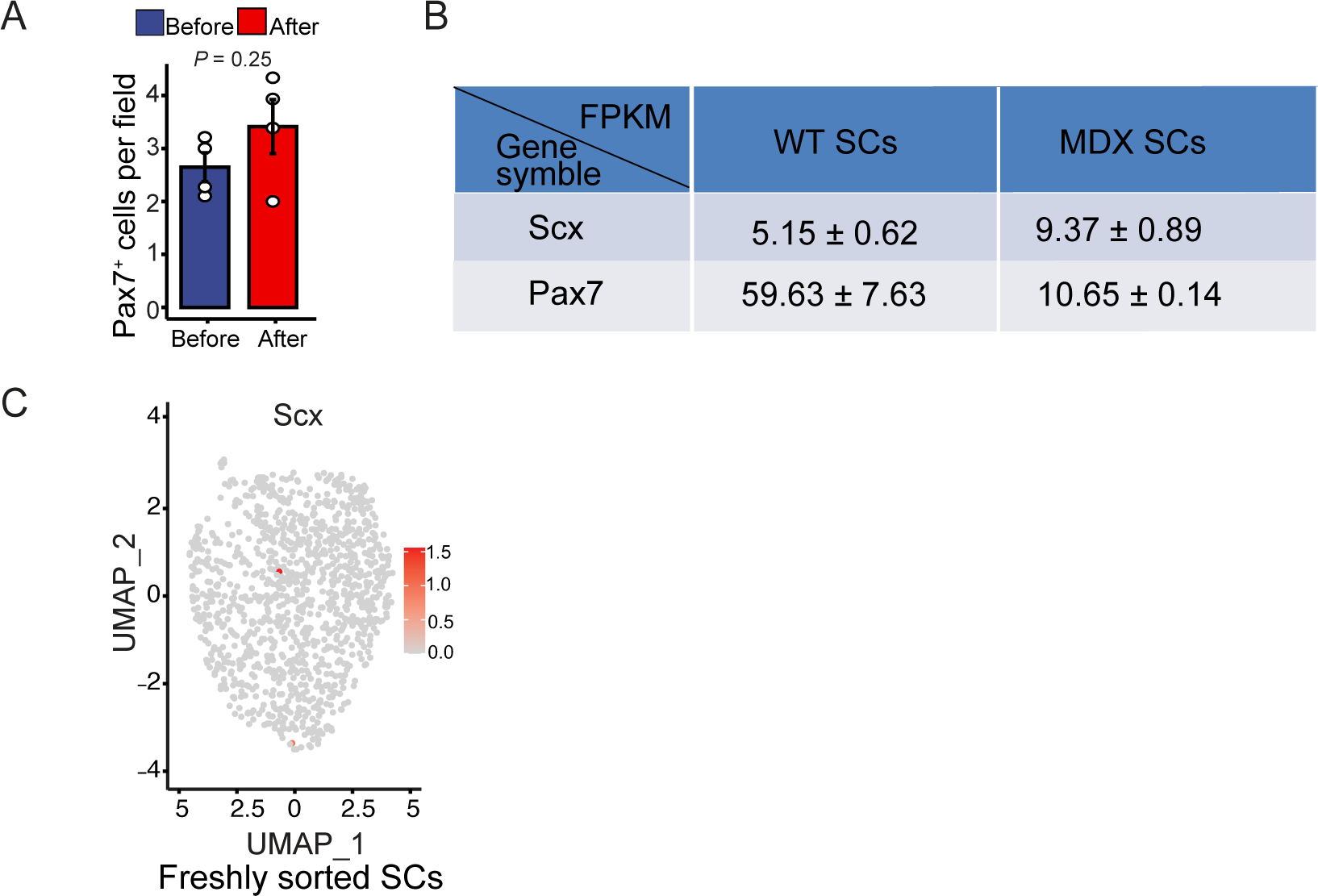
Additional information for Figure 2. A. Averaged Pax7^+^ SC number per field (0.06mm^2^) from data in B. Mouse and cell numbers are the same as in Figure 2C. B. Gene expression levels (in FPKM) of *Scx* and *Pax7* in SCs isolated from wild type (WT) and *mdx* mice using bulk-RNA-seq. Data are extracted from published data sets(Li, Rozo et al. 2019, Madaro, Torcinaro et al. 2019). C. UMAP plot of re-analyzed scRNA-seq for Scx expression in SCs isolated from uninjured muscles (freshly sorted) using published data in (Dell’Orso, Juan et al. 2019). Data are presented with the mean ± s.d.; the *P* value is indicated. Unpaired two-tailed Student’s *t*-test was applied.

**Figure S3:**
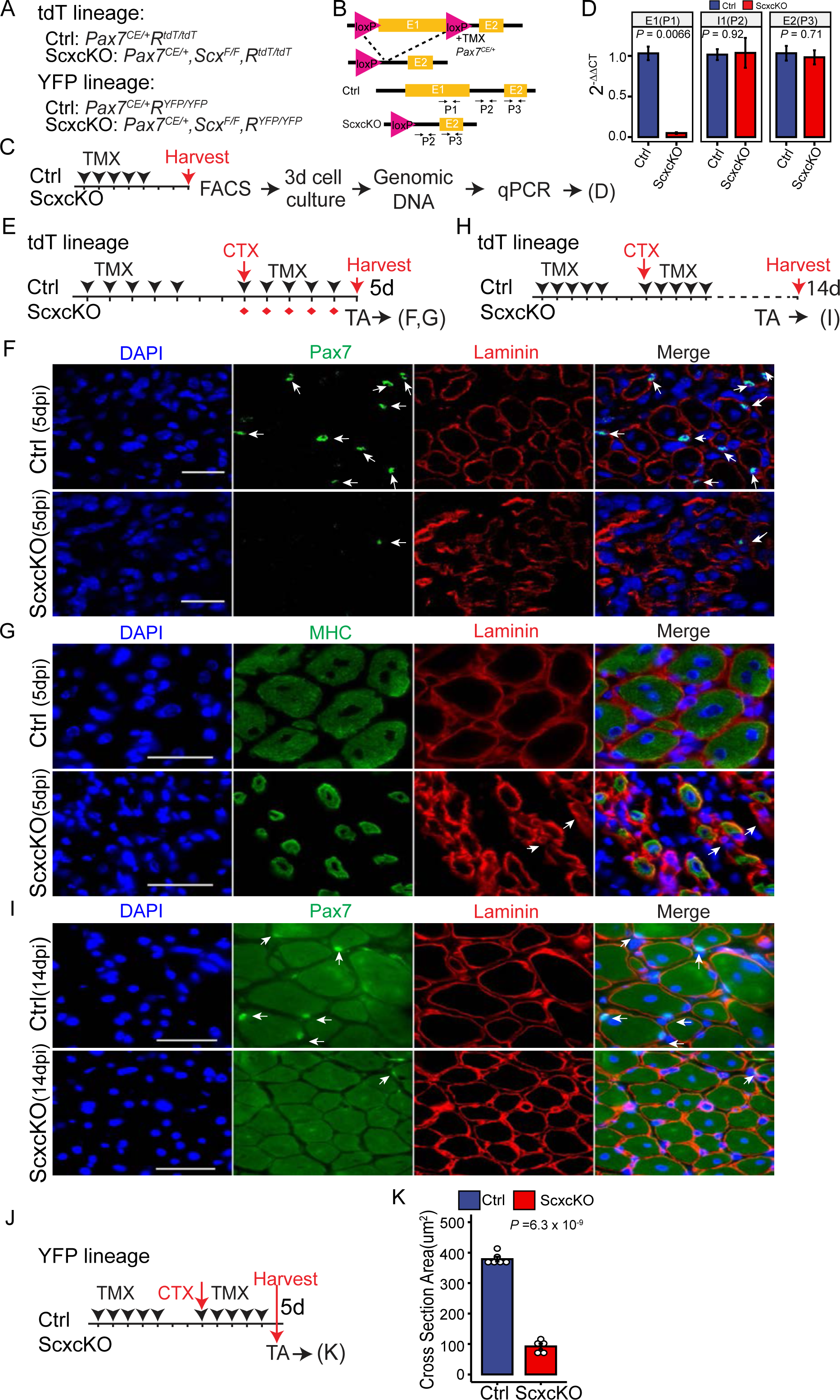
Addition data for Figure 3. A. Detailed description for the genotypes used as Ctrl and ScxcKO mice in Fig. 3. For tdT lineage, *R^tdT^* reporter was included. For YFP lineage, *R^YFP^* reporter was included. B. Depiction of *Scx* gene inactivation by TMX-induced recombination of loxP sites flanking the exon 1 of *Scx* using the *Pax7^CE^* allele. PCR primer sets P1, P2, and P3 were used to detect exon 1 (E1), intron 1 (I1) and E2 of the *Scx* gene, respectively. Primer sequences are in Table S10. C. Experimental scheme to determine the recombination efficiency using FACS-isolated Ctrl and ScxcKO SCs. Freshly sorted SCs were plated down and cultured in growth medium for 3 days and harvested for genomic DNA extraction and qPCR for data in (D). D. Relative levels of E1, I1, and E2 in control and ScxcKO myoblasts determined by qPCR, followed by 2^-ΔΔCt^ analysis. E. Experimental scheme as Fig. 3A for 5 dpi data in F-G. F-G. TA muscle sections (from E) were sectioned and stained with Pax7 and Laminin in (F), and MHC and laminin in (G). Arrows indicate Pax7^+^ SCs in (F), whereas arrows indicate Laminin^+^MHC^-^ ghost fibers (G) (N = 5 mice per group; n = 1748 control and n = 442 ScxcKO Pax7^+^ SCs). H. Experimental scheme as Fig. 3A for 14 dpi data in (I). I. TA muscle section from (H) from were sectioned and stained with Pax7 and Laminin (N = 5 mice per group; n = 350 control and n = 186 ScxcKO Pax7^+^ SCs examined). J. Same experimental scheme as in (E), except that *R^YFP^* reporter (YFP lineage), instead of *R^tdT^* reporter, was included. K. Histogram of the fiber CSA from 5 dpi TA muscle sections from (J). (N = 6 mice for control, and N = 5 mice for ScxcKO group) Nuclei were stained with DAPI. Scale bar = 20 μm. Data are presented with the mean ± s.d.; adjusted *P* values are shown. (D, K) Unpaired two-tailed Student’s *t*-test.

**Figure S4:**
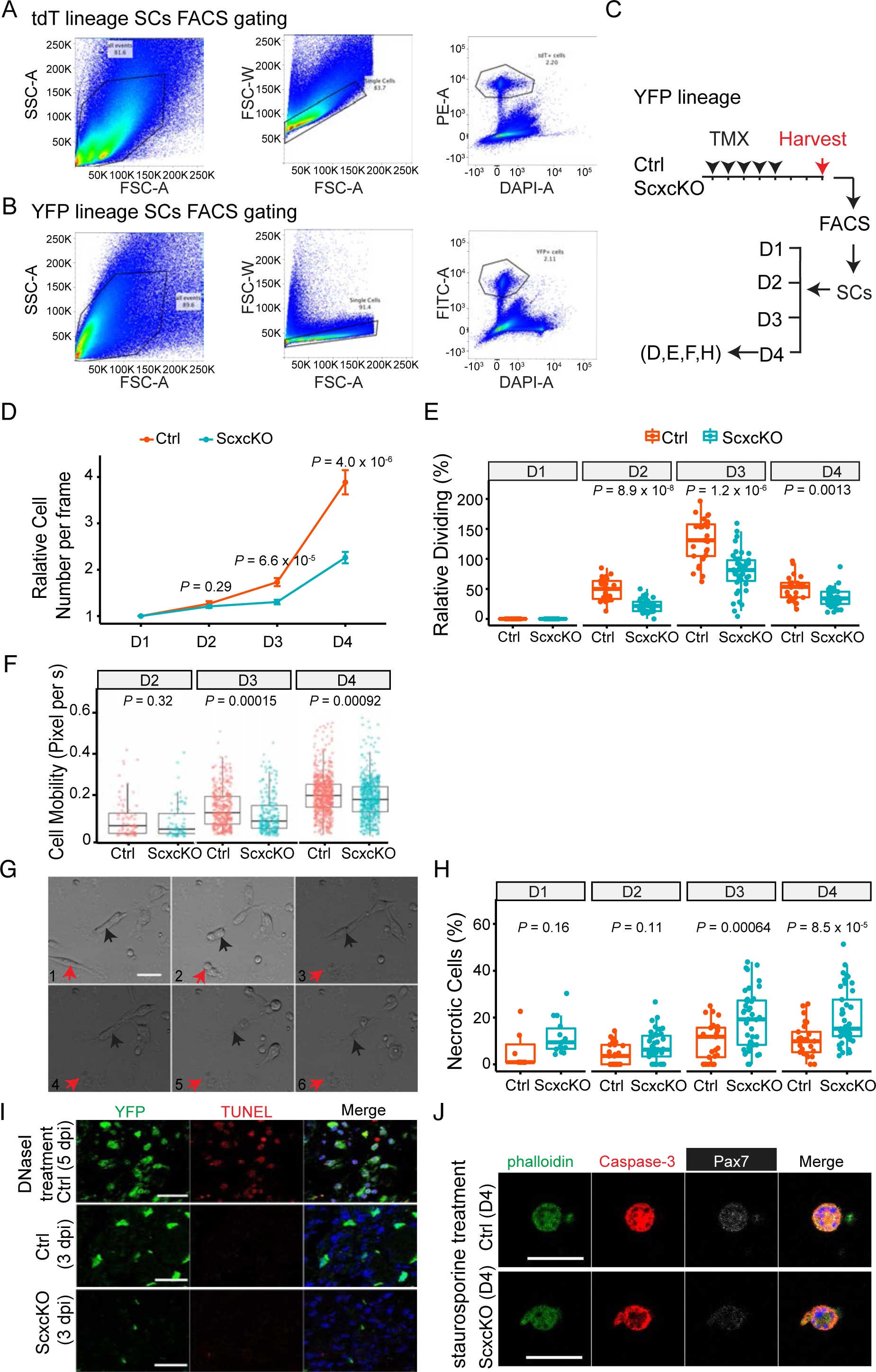
Additional data for Figure 4. A-B. FACS profiles for isolating tdT (A) and YFP (B) linage marked SCs. C. Similar experimental scheme as in Fig. 4 A for refence to live imaging data in (D-F). D. Line plot of relative cell number per frame; normalized to cell number of the Ctrl group at the beginning as 1. E. Box plot of relative dividing cell ratio. Relative dividing ratio is defined as the divided cell number of each day per group / the beginning cell number of that day. There were no dividing cells detected at D1, each dot represents one image data, 3 well per group. F. Box plot of cell migration speed based on tracking cell displacement by pixel (0.33 μm / pixel), each dot represents a cell data. G. Still frames from time lapse show examples of necrotic cells at D3 of the ScxcKO group. Frames 1 - 6 are sequential time lapse images (covering 500 mins). Two cells of focus are indicated by red and black arrowheads to track their appearance from a healthy state (earlier frames) to a necrotic state (later frames). H. Box plot of necrotic cell ratio. Necrotic cell was manually identified by morphology in each frame on each day, and normalized to the beginning cell number of that day, each dot represents one image data. I. For positive control, 5 dpi TA muscle from a *Pax7 ^CE/+^;Scx^Ty1/Ty1^;R^YFP/YFP^* mouse was treated with 200ugml^-1^ DNase I for 10 mins, followed by TUNEL assay and YFP staining (top panel; 4 sections examined and 5 images taken). Lower two panels are Ctrl and ScxcKO 3 dpi TA muscle sections subjected to TUNEL assay and YFP staining (N = 4 mice per group, 9 sections per slides per group, and 3-7 images per section analyzed). No appreciable number of TUNEL^+^YFP^+^ cells were found in Ctrl and ScxcKO, hence quantification omitted. J. FACS-isolated Ctrl and ScxcKO SCs were cultured in growth medium for 4 days. For positive control, 1 µM of staurosporine was added for 6 h prior to harvesting. Cells were then stained for cleaved Caspase-3 and Pax7; Phalloidin was used to identify cells by actin cytoskeleton. Nuclei were stained with DAPI; Scale bar = 20 μm. Data are presented with mean ± s.d.; adjusted *P* values are shown. Unpaired two-tailed Student’s *t*-tests were applied to data collected in each day.

**Figure S5:**
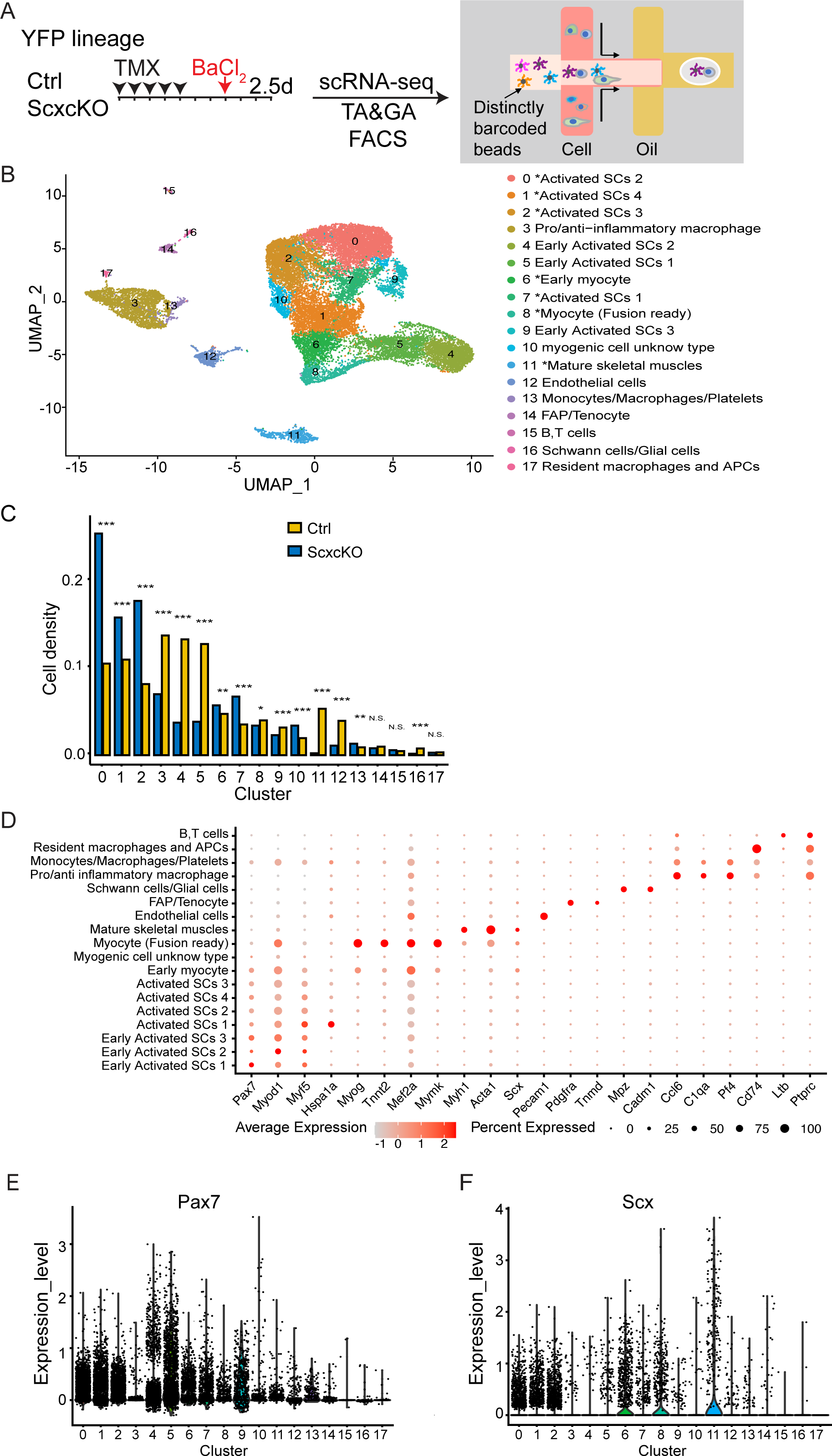
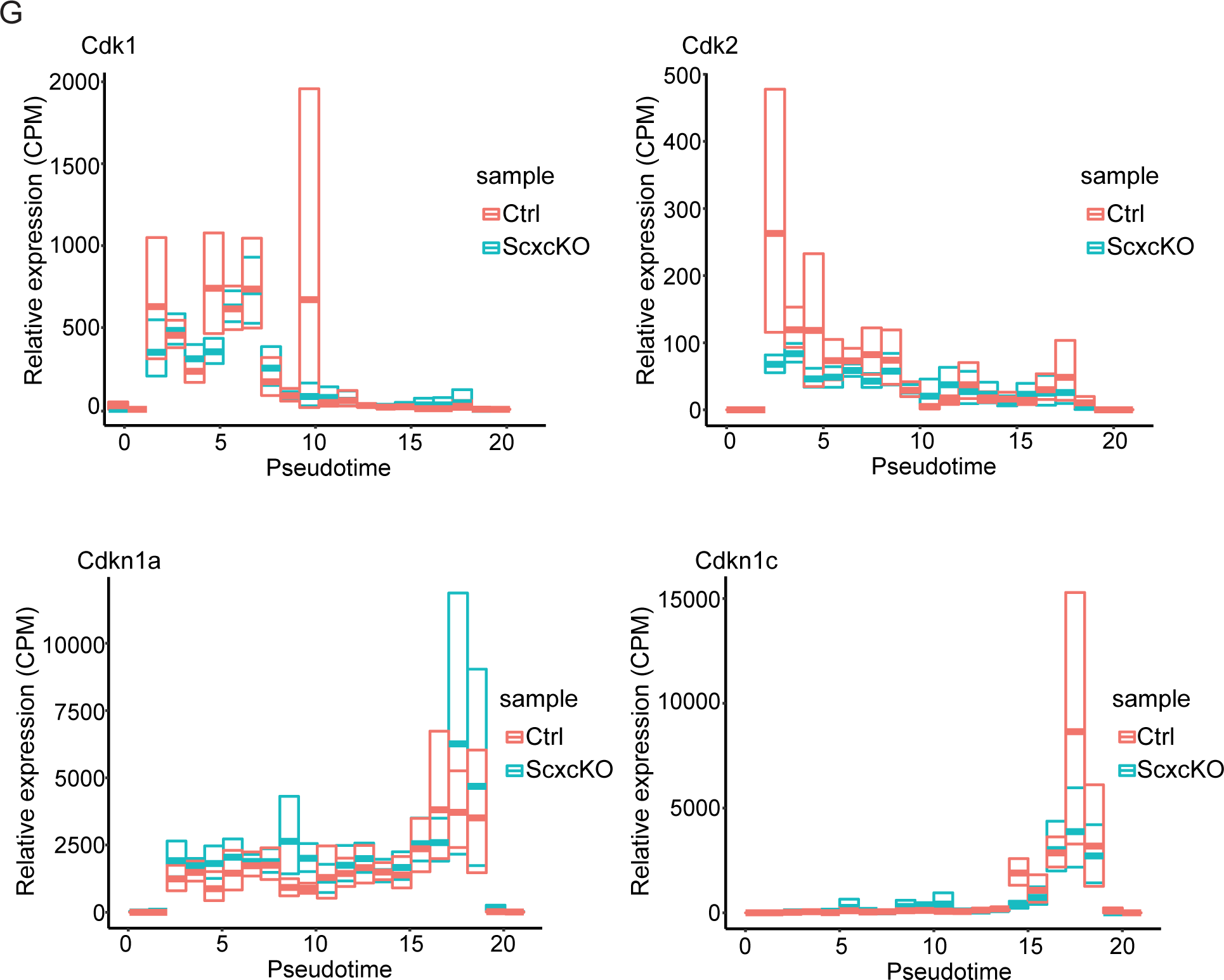
Additional analyses of data in Figure 5. A. Detailed scRNA-seq procedures diagram as in Fig. 5A for reference to analyses in (B-G). B. UMAP plot of combined scRNA-seq of Ctrl and ScxcKO cells. A total of 11,388 control cells and 12,844 ScxcKO cells were included for analysis, and 18 cell clusters were delineated (shown by different colors and with assigned cell types indicated to the right). Asterisks denote myogenic clusters subjected to further analyses in Figure 5B, C, H, I. C. Cell density distribution of each cell cluster in Ctrl and ScxcKO samples. NS, not significant; \**P* < 0.05; \*\**P* < 0.01; \*\*\**P* < 0.001. D. Expression of select marker genes in each cell cluster. Dot size represents the percentage of expressed cells within each cluster, whereas color intensity represents relative expression level (keys at bottom). E. *Pax7* expression levels and cell numbers in each cluster; each dot represents one cell. F. *Scx* expression levels and cell numbers in each cluster. G. Relative expression levels (CPM, counts per million UMI) of 4 select cell cycle genes along the pseudotime depicted in Figure 5C.

**Figure S6:**
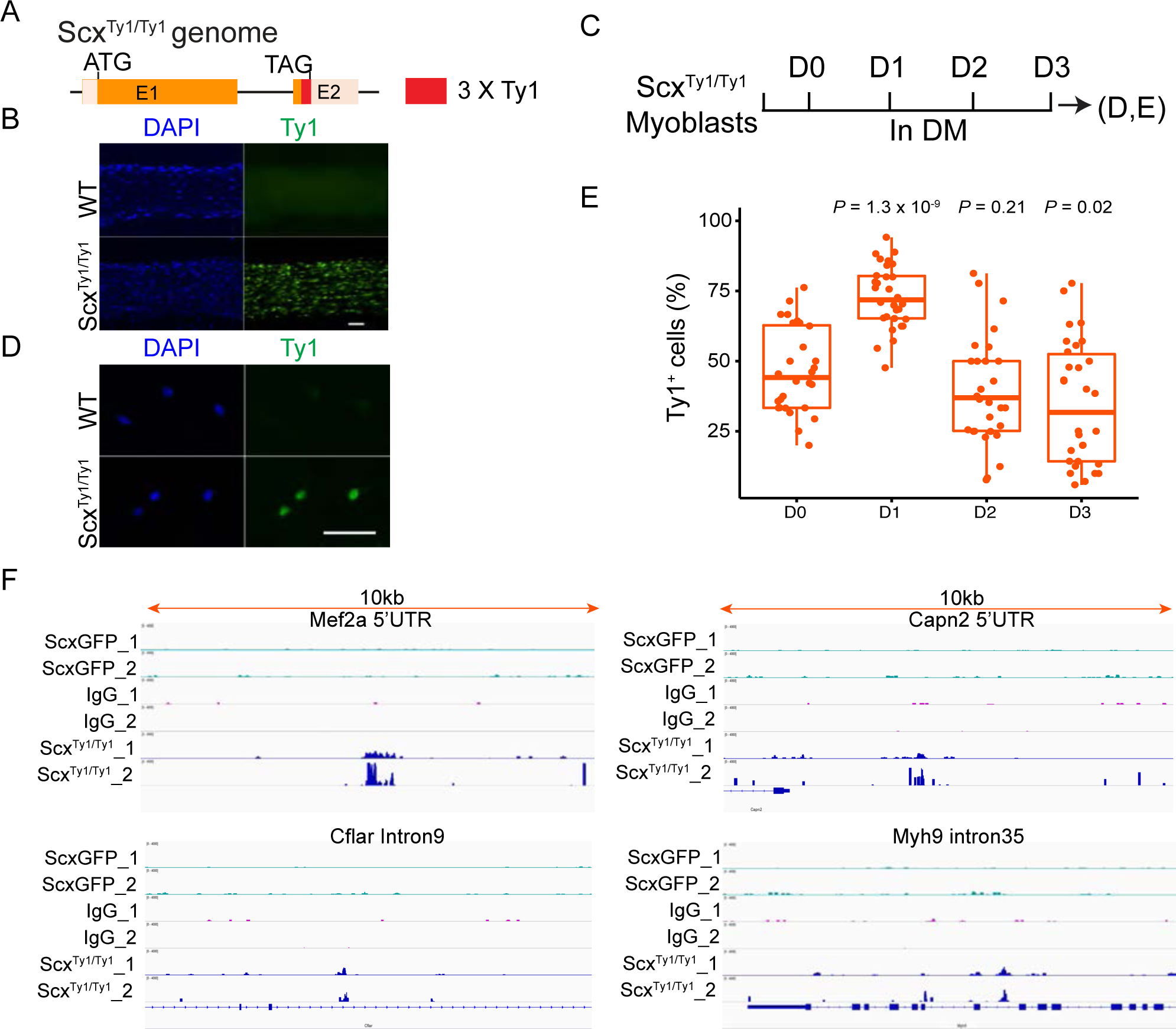
Additional data to support Figure 6. A. Diagram of the genomic structure of *Scx^Ty1^* allele. Three (3X) Ty1 tags were inserted just before the TGA codon of the *Scx* gene. B. Neonatal patellar tendons of Scx^Ty1/Ty1^ and wild type (WT) mice were fixed, sectioned, and stained for Ty1. C. *In vitro* myogenic differentiation scheme using Scx^Ty1/Ty1^ SC-derived myoblasts. Myoblasts were cultured in GM for 12 h (D0), and switched to DM for 3 days. Cells in each day were stained for Ty1 (D0-3). D. Scx^Ty1/Ty1^ and WT myoblasts at D0 were stained for Ty1 as an example for data in (E). E. Boxplot of percentages of cells with detectable Ty1 signal assayed at different time points listed in (C). Each dot represents one image data, 3 wells per time point, 10 images per well, and a total of 2785 cells examined. N = 3 mice. Data are present as the mean ± s.d.; adjusted *P* values are shown. Unpaired two-tailed Student’s *t*-tests were applied, D0 sample as the refence group. F, Genomic snapshots of Scx CUT&RUN signals across a 10 Kb region of each of the 4 select genes in Figure 6G. The green lines represent CUT&RUN signal from anti-Ty1 on ScxGFP myoblasts; the magenta and blue lines represent the IgG control and anti-Ty1 antibody CUT&RUN signals detected in Scx^Ty1/Ty1^ primary myoblasts, respectively. Nuclei were stained with DAPI; Scale bar = 20 μm.

## Notes

https://figshare.com/projects/Skeletal_Muscle_Satellite_Cells_Co-Opt_the_Tenogenic_Gene_Scleraxis_to_Instruct_Regeneration/190935

### Competing Interest Statement

The authors have declared no competing interest.

### Summary of Updates

new plot was added or modified to F2E, F4F-G, SF2C, and SF4I-J, figure legends and the manuscript has been revised according to the related figures.

## References

Blondelle, J., Y. Ohno, V. Gache, S. Guyot, S. Storck, N. Blanchard-Gutton, I. Barthelemy, G. Walmsley, A. Rahier, S. Gadin, M. Maurer, L. Guillaud, A. Prola, A. Ferry, G. Aubin-Houzelstein, J. Demarquoy, F. Relaix, R. J. Piercy, S. Blot, A. Kihara, L. Tiret and F. Pilot-Storck (2015). “HACD1, a regulator of membrane composition and fluidity, promotes myoblast fusion and skeletal muscle growth.” J Mol Cell Biol 7(5): 429–440.

Brent, A. E., R. Schweitzer and C. J. Tabin (2003). “A somitic compartment of tendon progenitors.” Cell 113(2): 235–248.

Budd, R. C., W. C. Yeh and J. Tschopp (2006). “cFLIP regulation of lymphocyte activation and development.” Nat Rev Immunol 6(3): 196–204.

Cai, H., F. Guo, S. Wen, X. Jin, H. Wu and D. Ren (2022). “Overexpressed integrin alpha 2 inhibits the activation of the transforming growth factor beta pathway in pancreatic cancer via the TFCP2-SMAD2 axis.” J Exp Clin Cancer Res 41(1): 73.

Chuang, Y. C., H. Y. Wu, Y. L. Lin, S. C. Tzou, C. H. Chuang, T. Y. Jian, P. R. Chen, Y. C. Chang, C. H. Lin, T. H. Huang, C. C. Wang, Y. L. Chan and K. W. Liao (2018). “Blockade of ITGA2 Induces Apoptosis and Inhibits Cell Migration in Gastric Cancer.” Biol Proced Online 20: 10.

Cserjesi, P. (1995). “Scleraxis: a basic helix-loop-helix protein that prefigures skeletal formation during mouse embryogenesis.” Development 121: 1099–1110

De Micheli, A. J., E. J. Laurilliard, C. L. Heinke, H. Ravichandran, P. Fraczek, S. Soueid-Baumgarten, I. De Vlaminck, O. Elemento and B. D. Cosgrove (2020). “Single-Cell Analysis of the Muscle Stem Cell Hierarchy Identifies Heterotypic Communication Signals Involved in Skeletal Muscle Regeneration.” Cell Rep 30(10): 3583–3595 e3585.

Dedieu, S., G. Mazeres, N. Dourdin, P. Cottin and J. J. Brustis (2003). “Transactivation of capn2 by myogenic regulatory factors during myogenesis.” J Mol Biol 326(2): 453–465.

Dell’Orso, S., A. H. Juan, K. D. Ko, F. Naz, J. Perovanovic, G. Gutierrez-Cruz, X. Feng and V. Sartorelli (2019). “Single cell analysis of adult mouse skeletal muscle stem cells in homeostatic and regenerative conditions.” Development 146(12).

Doherty, K. R., A. R. Demonbreun, G. Q. Wallace, A. Cave, A. D. Posey, K. Heretis, P. Pytel and E. M. McNally (2008). “The endocytic recycling protein EHD2 interacts with myoferlin to regulate myoblast fusion.” J Biol Chem 283(29): 20252–20260.

Dubinska-Magiera, M., M. Zaremba-Czogalla and R. Rzepecki (2013). “Muscle development, regeneration and laminopathies: how lamins or lamina-associated proteins can contribute to muscle development, regeneration and disease.” Cell Mol Life Sci 70(15): 2713–2741.

Esteves de Lima, J., C. Blavet, M. A. Bonnin, E. Hirsinger, G. Comai, L. Yvernogeau, M. C. Delfini, L. Bellenger, S. Mella, S. Nassari, C. Robin, R. Schweitzer, C. Fournier-Thibault, T. Jaffredo, S. Tajbakhsh, F. Relaix and D. Duprez (2021). “Unexpected contribution of fibroblasts to muscle lineage as a mechanism for limb muscle patterning.” Nat Commun 12(1): 3851.

Estrella, N. L., C. A. Desjardins, S. E. Nocco, A. L. Clark, Y. Maksimenko and F. J. Naya (2015). “MEF2 transcription factors regulate distinct gene programs in mammalian skeletal muscle differentiation.” J Biol Chem 290(2): 1256–1268.

Ewels, P. A., A. Peltzer, S. Fillinger, H. Patel, J. Alneberg, A. Wilm, M. U. Garcia, P. Di Tommaso and S. Nahnsen (2020). “The nf-core framework for community-curated bioinformatics pipelines.” Nat Biotechnol 38(3): 276–278.

Francetic T, L. Q. (2011). “Skeletal myogenesis and Myf5 activation.”

Fukada, S. I., T. Higashimoto and A. Kaneshige (2022). “Differences in muscle satellite cell dynamics during muscle hypertrophy and regeneration.” Skelet Muscle 12(1): 17.

Giordani, L., G. J. He, E. Negroni, H. Sakai, J. Y. C. Law, M. M. Siu, R. Wan, A. Corneau, S. Tajbakhsh, T. H. Cheung and F. Le Grand (2019). “High-Dimensional Single-Cell Cartography Reveals Novel Skeletal Muscle-Resident Cell Populations.” Mol Cell 74(3): 609–621 e606.

Gumucio, J. P., M. M. Schonk, Y. A. Kharaz, E. Comerford and C. L. Mendias (2020). “Scleraxis is required for the growth of adult tendons in response to mechanical loading.” JCI Insight 5(13).

Hao, Y., S. Hao, E. Andersen-Nissen, W. M. Mauck, 3rd, S. Zheng, A. Butler, M. J. Lee, A. J. Wilk, C. Darby, M. Zager, P. Hoffman, M. Stoeckius, E. Papalexi, E. P. Mimitou, J. Jain, A. Srivastava, T. Stuart, L. M. Fleming, B. Yeung, A. J. Rogers, J. M. McElrath, C. A. Blish, R. Gottardo, P. Smibert and R. Satija (2021). “Integrated analysis of multimodal single-cell data.” Cell 184(13): 3573-3587 e3529.

Harvey, T., S. Flamenco and C. M. Fan (2019). “A Tppp3(+)Pdgfra(+) tendon stem cell population contributes to regeneration and reveals a shared role for PDGF signalling in regeneration and fibrosis.” Nat Cell Biol 21(12): 1490–1503.

Hernandez-Hernandez, J. M., E. G. Garcia-Gonzalez, C. E. Brun and M. A. Rudnicki (2017). “The myogenic regulatory factors, determinants of muscle development, cell identity and regeneration.” Semin Cell Dev Biol 72: 10–18.

Honda, M., F. Masui, N. Kanzawa, T. Tsuchiya and T. Toyo-oka (2008). “Specific knockdown of m-calpain blocks myogenesis with cDNA deduced from the corresponding RNAi.” Am J Physiol Cell Physiol 294(4): C957–965.

Howell, K., C. Chien, R. Bell, D. Laudier, S. F. Tufa, D. R. Keene, N. Andarawis-Puri and A. H. Huang (2017). “Novel Model of Tendon Regeneration Reveals Distinct Cell Mechanisms Underlying Regenerative and Fibrotic Tendon Healing.” Sci Rep 7: 45238.

Huang, Y., F. Clarke, M. Karimi, N. H. Roy, E. K. Williamson, M. Okumura, K. Mochizuki, E. J. Chen, T. J. Park, G. F. Debes, Y. Zhang, T. Curran, T. Kambayashi and J. K. Burkhardt (2015). “CRK proteins selectively regulate T cell migration into inflamed tissues.” J Clin Invest 125(3): 1019–1032.

Kardon, G. (1998). “Muscle and tendon morphogenesis in the avian hind limb.” Development 125(20): 4019–4032.

Kaya-Okur, H. S., D. H. Janssens, J. G. Henikoff, K. Ahmad and S. Henikoff (2020). “Efficient low-cost chromatin profiling with CUT&Tag.” Nat Protoc 15(10): 3264–3283.

Korcari, A., S. Muscat, E. McGinn, M. R. Buckley and A. E. Loiselle (2022). “Depletion of Scleraxis-lineage cells during tendon healing transiently impairs multi-scale restoration of tendon structure during early healing.” PLoS One 17(10): e0274227.

Lepper, C., S. J. Conway and C. M. Fan (2009). “Adult satellite cells and embryonic muscle progenitors have distinct genetic requirements.” Nature 460(7255): 627–631.

Li, L., M. Rozo, S. Yue, X. Zheng, J. T. F, C. Lepper and C. M. Fan (2019). “Muscle stem cell renewal suppressed by Gas1 can be reversed by GDNF in mice.” Nat Metab 1(10): 985–995.

Li, Y., T. Wu and S. Liu (2021). “Identification and Distinction of Tenocytes and Tendon-Derived Stem Cells.” Front Cell Dev Biol 9: 629515.

Lin, X., X. Yang, Q. Li, Y. Ma, S. Cui, D. He, X. Lin, R. J. Schwartz and J. Chang (2012). “Protein tyrosine phosphatase-like A regulates myoblast proliferation and differentiation through MyoG and the cell cycling signaling pathway.” Mol Cell Biol 32(2): 297–308.

Liu, L., T. H. Cheung, G. W. Charville and T. A. Rando (2015). “Isolation of skeletal muscle stem cells by fluorescence-activated cell sorting.” Nat Protoc 10(10): 1612–1624.

Liu, N., B. R. Nelson, S. Bezprozvannaya, J. M. Shelton, J. A. Richardson, R. Bassel-Duby and E. N. Olson (2014). “Requirement of MEF2A, C, and D for skeletal muscle regeneration.” Proc Natl Acad Sci U S A 111(11): 4109–4114.

Madaro, L., A. Torcinaro, M. De Bardi, F. F. Contino, M. Pelizzola, G. R. Diaferia, G. Imeneo, M. Bouche, P. L. Puri and F. De Santa (2019). “Macrophages fine tune satellite cell fate in dystrophic skeletal muscle of mdx mice.” PLoS Genet 15(10): e1008408.

Madisen, L., T. A. Zwingman, S. M. Sunkin, S. W. Oh, H. A. Zariwala, H. Gu, L. L. Ng, R. D. Palmiter, M. J. Hawrylycz, A. R. Jones, E. S. Lein and H. Zeng (2010). “A robust and high-throughput Cre reporting and characterization system for the whole mouse brain.” Nat Neurosci 13(1): 133–140.

Maggi, L., N. Carboni and P. Bernasconi (2016). “Skeletal Muscle Laminopathies: A Review of Clinical and Molecular Features.” Cells 5(3).

McGinnis, C. S., L. M. Murrow and Z. J. Gartner (2019). “DoubletFinder: Doublet Detection in Single-Cell RNA Sequencing Data Using Artificial Nearest Neighbors.” Cell Syst 8(4): 329–337 e324.

Morton, A. B., C. E. Norton, N. L. Jacobsen, C. A. Fernando, D. D. W. Cornelison and S. S. Segal (2019). “Barium chloride injures myofibers through calcium-induced proteolysis with fragmentation of motor nerves and microvessels.” Skelet Muscle 9(1): 27.

Muhl, L., G. Genove, S. Leptidis, J. Liu, L. He, G. Mocci, Y. Sun, S. Gustafsson, B. Buyandelger, I. V. Chivukula, A. Segerstolpe, E. Raschperger, E. M. Hansson, J. L. M. Bjorkegren, X. R. Peng, M. Vanlandewijck, U. Lendahl and C. Betsholtz (2020). “Single-cell analysis uncovers fibroblast heterogeneity and criteria for fibroblast and mural cell identification and discrimination.” Nat Commun 11(1): 3953.

Murach, K. A., B. D. Peck, R. A. Policastro, I. J. Vechetti, D. W. Van Pelt, C. M. Dungan, L. T. Denes, X. Fu, C. R. Brightwell, G. E. Zentner, E. E. Dupont-Versteegden, C. I. Richards, J. J. Smith, C. S. Fry, J. J. McCarthy and C. A. Peterson (2021). “Early satellite cell communication creates a permissive environment for long-term muscle growth.” iScience 24(4): 102372.

Murchison, N. D., B. A. Price, D. A. Conner, D. R. Keene, E. N. Olson, C. J. Tabin and R. Schweitzer (2007). “Regulation of tendon differentiation by scleraxis distinguishes force-transmitting tendons from muscle-anchoring tendons.” Development 134(14): 2697–2708.

Nguyen, Q. G., J. N. Buskin, C. L. Himeda, M. A. Shield and S. D. Hauschka (2003). “Differences in the function of three conserved E-boxes of the muscle creatine kinase gene in cultured myocytes and in transgenic mouse skeletal and cardiac muscle.” J Biol Chem 278(47): 46494–46505.

Ono, Y., S. Schlesinger, K. Fukunaga, S. Yambe, T. Sato, T. Sasaki, C. Shukunami, H. Asahara and M. Inui (2023). “Scleraxis-lineage cells are required for correct muscle patterning.” Development 150(10).

Oprescu, S. N., F. Yue, J. Qiu, L. F. Brito and S. Kuang (2020). “Temporal Dynamics and Heterogeneity of Cell Populations during Skeletal Muscle Regeneration.” iScience 23(4): 100993.

Paxton, C. W., R. A. Cosgrove, A. C. Drozd, E. L. Wiggins, S. Woodhouse, R. A. Watson, H. J. Spence, B. W. Ozanne and J. M. Pell (2011). “BTB-Kelch protein Krp1 regulates proliferation and differentiation of myoblasts.” Am J Physiol Cell Physiol 300(6): C1345–1355.

Posey, A. D., Jr., P. Pytel, K. Gardikiotes, A. R. Demonbreun, M. Rainey, M. George, H. Band and E. M. McNally (2011). “Endocytic recycling proteins EHD1 and EHD2 interact with fer-1-like-5 (Fer1L5) and mediate myoblast fusion.” J Biol Chem 286(9): 7379-7388.

Pryce, B. A., A. E. Brent, N. D. Murchison, C. J. Tabin and R. Schweitzer (2007). “Generation of transgenic tendon reporters, ScxGFP and ScxAP, using regulatory elements of the scleraxis gene.” Dev Dyn 236(6): 1677–1682.

Qiu, X., Q. Mao, Y. Tang, L. Wang, R. Chawla, H. A. Pliner and C. Trapnell (2017). “Reversed graph embedding resolves complex single-cell trajectories.” Nat Methods 14(10): 979–982.

Ramirez-Martinez, A., B. K. Cenik, S. Bezprozvannaya, B. Chen, R. Bassel-Duby, N. Liu and E. N. Olson (2017). “KLHL41 stabilizes skeletal muscle sarcomeres by nonproteolytic ubiquitination.” Elife 6.

Ren, D., J. Zhao, Y. Sun, D. Li, Z. Meng, B. Wang, P. Fan, Z. Liu, X. Jin and H. Wu (2019). “Overexpressed ITGA2 promotes malignant tumor aggression by up-regulating PD-L1 expression through the activation of the STAT3 signaling pathway.” J Exp Clin Cancer Res 38(1): 485.

S Srinivas 1, T. W., C S Lin, C M William, Y Tanabe, T M Jessell, F Costantini (2001). “Cre reporter strains produced by targeted insertion of EYFP and ECFP into the ROSA26 locus.” BMC Developmental Biology.

Sakabe, T., K. Sakai, T. Maeda, A. Sunaga, N. Furuta, R. Schweitzer, T. Sasaki and T. Sakai (2018). “Transcription factor scleraxis vitally contributes to progenitor lineage direction in wound healing of adult tendon in mice.” J Biol Chem 293(16): 5766–5780.

Schweitzer, R., J. H. Chyung, L. C. Murtaugh, A. E. Brent, V. Rosen, E. N. Olson, A. Lassar and C. J. Tabin (2001). “Analysis of the tendon cell fate using Scleraxis, a specific marker for tendons and ligaments.” Development 128(19): 3855–3866.

Scott, R. W., M. Arostegui, R. Schweitzer, F. M. V. Rossi and T. M. Underhill (2019). “Hic1 Defines Quiescent Mesenchymal Progenitor Subpopulations with Distinct Functions and Fates in Skeletal Muscle Regeneration.” Cell Stem Cell 25(6): 797–813 e799.

Senf, S. M. (2013). “Skeletal muscle heat shock protein 70: diverse functions and therapeutic potential for wasting disorders.” Front Physiol 4: 330.

Seok, H. Y., M. Tatsuguchi, T. E. Callis, A. He, W. T. Pu and D. Z. Wang (2011). “miR-155 inhibits expression of the MEF2A protein to repress skeletal muscle differentiation.” J Biol Chem 286(41): 35339–35346.

Shukunami, C., A. Takimoto, Y. Nishizaki, Y. Yoshimoto, S. Tanaka, S. Miura, H. Watanabe, T. Sakuma, T. Yamamoto, G. Kondoh and Y. Hiraki (2018). “Scleraxis is a transcriptional activator that regulates the expression of Tenomodulin, a marker of mature tenocytes and ligamentocytes.” Sci Rep 8(1): 3155.

Shukunami, C., A. Takimoto, M. Oro and Y. Hiraki (2006). “Scleraxis positively regulates the expression of tenomodulin, a differentiation marker of tenocytes.” Dev Biol 298(1): 234–247.

Strenzke, M., P. Alberton, A. Aszodi, D. Docheva, E. Haas, C. Kammerlander, W. Bocker and M. M. Saller (2020). “Tenogenic Contribution to Skeletal Muscle Regeneration: The Secretome of Scleraxis Overexpressing Mesenchymal Stem Cells Enhances Myogenic Differentiation In Vitro.” Int J Mol Sci 21(6).

Swailes, N. T., M. Colegrave, P. J. Knight and M. Peckham (2006). “Non-muscle myosins 2A and 2B drive changes in cell morphology that occur as myoblasts align and fuse.” J Cell Sci 119(Pt 17): 3561–3570.

Tozer, S. and D. Duprez (2005). “Tendon and ligament: development, repair and disease.” Birth Defects Res C Embryo Today 75(3): 226–236.

Vesely, E. D., C. W. Heilig and F. C. Brosius, 3rd (2009). “GLUT1-induced cFLIP expression promotes proliferation and prevents apoptosis in vascular smooth muscle cells.” Am J Physiol Cell Physiol **297**(3): C759-765.

Wales, S., S. Hashemi, A. Blais and J. C. McDermott (2014). “Global MEF2 target gene analysis in cardiac and skeletal muscle reveals novel regulation of DUSP6 by p38MAPK-MEF2 signaling.” Nucleic Acids Res 42(18): 11349–11362.

Wang, W., C. Z. Prince, Y. Mou and M. J. Pollman (2002). “Notch3 signaling in vascular smooth muscle cells induces c-FLIP expression via ERK/MAPK activation. Resistance to Fas ligand-induced apoptosis.” J Biol Chem 277(24): 21723–21729.

Wang, Y. N., W. C. Yang, P. W. Li, H. B. Wang, Y. Y. Zhang and L. S. Zan (2018). “Myocyte enhancer factor 2A promotes proliferation and its inhibition attenuates myogenic differentiation via myozenin 2 in bovine skeletal muscle myoblast.” PLoS One 13(4): e0196255.

Webster, M. T., U. Manor, J. Lippincott-Schwartz and C. M. Fan (2016). “Intravital Imaging Reveals Ghost Fibers as Architectural Units Guiding Myogenic Progenitors during Regeneration.” Cell Stem Cell 18(2): 243–252.

Yaseen, W., O. Kraft-Sheleg, S. Zaffryar-Eilot, S. Melamed, C. Sun, D. P. Millay and P. Hasson (2021). “Fibroblast fusion to the muscle fiber regulates myotendinous junction formation.” Nat Commun 12(1): 3852.

Yin, H., F. Price and M. A. Rudnicki (2013). “Satellite cells and the muscle stem cell niche.” Physiol Rev 93(1): 23–67.

Yoshimoto, Y., A. Takimoto, H. Watanabe, Y. Hiraki, G. Kondoh and C. Shukunami (2017). “Scleraxis is required for maturation of tissue domains for proper integration of the musculoskeletal system.” Sci Rep 7: 45010.

Yue, L., R. Wan, S. Luan, W. Zeng and T. H. Cheung (2020). “Dek Modulates Global Intron Retention during Muscle Stem Cells Quiescence Exit.” Dev Cell 53(6): 661–676 e666.

Zhang, N. and Y. W. He (2005). “An essential role for c-FLIP in the efficient development of mature T lymphocytes.” J Exp Med 202(3): 395–404.

